# Characterizing Material Mechanics and Modeling Variability in Organic Hydrogels: A Framework for Soft and Biodegradable Robotics

**DOI:** 10.64898/2026.11.25.769239

**Authors:** Avery Sonora Williamson, Michael J. Bennington, Victoria A. Webster Wood

## Abstract

Organic hydrogels are increasingly used in biofabrication, soft robotics, and biomedical devices due to their tunable mechanical properties, biocompatibility, biodegradability, and flexible fabrication modalities, including embedded 3D printing. However, both polymer chemistry and fabrication conditions can introduce substantial variability in material behavior, particularly for biologically-derived polymers whose inconsistent molecular composition and cross-linking response affect viscoelastic performance. This variability complicates design, reduces reproducibility, and obscures the steady-state mechanical properties required for functional deployment. In this work, we present an experimental-computational framework for characterizing and predicting the mechanical behavior of 3D-printed hydrogels. The workflow characterizes material behavior across molecular, bulk, and population scales. First, we quantify intrinsic polymer variability via ^1^H-NMR spectroscopy to assess monomer distributions governing ionic cross-linking, establishing a molecular-level explanation for observed material variability. Next, we systematically measure how cross-linking time and temperature influence dynamic swelling and stiffness, highlighting representative conditions under which the material approaches steady-state mechanics while maintaining geometric fidelity. Finally, we develop a generative population model to describe geometry and stiffness evolution during cross-linking and calibrate its parameters using Sequential Monte Carlo (SMC) methods, enabling prediction of population-level distributions and variability from limited experimental data. We demonstrate this framework using FRESH-printed sodium alginate constructs as a case study; alginate hydrogels approach steady-state swelling and stiffness after extended cross-linking (>8 hr), and elevated temperatures increase contraction without improving modulus, indicating that low-temperature, long-duration cross-linking enhances geometric and mechanical stability. By integrating molecular composition, fabrication-dependent variability, and dynamic cross-linking behavior into a probabilistic model, this framework provides a structured approach for researchers to quantify uncertainty and design around variability in organic hydrogels, supporting more predictable deployment of soft robotic actuators, tissue-mimetic constructs, and biodegradable devices.

## Introduction

Soft robotic systems are increasingly used in biomedical devices, environmental monitoring, and human–robot interaction because their compliant architecture enables safe, adaptive engagement with sensitive biological tissues, organisms, and fragile ecosystems in ways that traditional rigid robots do not [1, 2]. Compliant actuators have been utilized in swimming [3–6] and crawling robots [7–9], grippers [10], and tentacles [11]. Recently, researchers have deployed robotic devices capable of locomotion in amphibious environments through hydraulic or electrical stimulation methods [6–8, 12–15]. However, many soft robotic systems still rely on synthetic elastomers and plastics that may persist or bioaccumulate when deployed in sensitive ecosystems or in the human body [16–18]. These limitations motivate the development of compliant robotic materials that are not only functional, but biologically and environmentally compatible.

Biologically sourced and biodegradable organic hydrogels offer a compelling alternative because they combine tunable mechanical properties, biocompatibility, and compatibility with aqueous environments. Their water-rich polymer networks allow stiffness, swelling, and viscoelasticity to be adjusted through polymer concentration, cross-linking chemistry, and ionic environment [19]. Among these materials, alginate, a naturally derived polysaccharide from seaweed, is particularly attractive due to its biodegradability, edibility for marine organisms, and ability to form ionically cross-linked networks with tunable stiffness and reversible shape-morphing capabilities [11, 19].

To leverage these properties in complex geometries, researchers increasingly use embedded printing approaches, in which soft or liquid-phase inks are extruded within a supportive medium to enable high-fidelity fabrication of compliant structures. This class of techniques, used across soft robotics and biofabrication [20], allows organic hydrogels to be patterned with a level of geometric freedom not achievable through traditional molding or casting. Within this broader family, Freeform Reversible Embedding of Suspended Hydrogels (FRESH) printing provides a versatile platform for printing biologically derived inks with fine feature resolution and minimal deformation [21]. Our previous demonstrations of bending and linear actuators, small-scale robotic grippers, and multi-actuator end-effectors fabricated via FRESH printing [11, 22] highlight the potential of alginate hydrogels for compliant, biodegradable robotic systems and devices.

Despite the demonstrated functional performance of organic hydrogels for soft robotics, material variability remains a major barrier to the reliable design and deployment of hydrogel-based actuators. Ionically cross-linked hydrogels depend strongly on diffusion[23], making the final mechanical properties sensitive to time, temperature, and ion concentration during fabrication [24, 25]. Additionally, the molecular composition of bio-derived materials can vary widely across natural sources. In alginate, the ratio of mannuronic (M) to guluronic (G) monomers can range from 0.21 to 1.59 in commonly harvested species [26], and this G/M ratio directly influences network formation, stiffness, and swelling behavior[19, 27, 28]. Similar variability is widespread across organic hydrogels, including gelatin, collagen, and decellularized extracellular matrix materials, where batch-to-batch differences in molecular weight distribution, cross-linking kinetics, and precursor purity similarly propagate into the bulk mechanical response [20, 29, 30]. In FRESH methods especially, the effects of fabrication-dependent factors such as ink viscosity, support-bath rheology, and peri-printing cross-linking conditions are inherently tied to polymer network behavior, making it difficult to isolate sources of variability between the material and manufacturing process. These molecular- and macro-level differences pose challenges for reproducibility and the prediction of actuator behavior, especially when typical industry sample sizes for isotropic elastomer materials (*N ≥* 3, ASTM D575/D412 [31, 32]) fail to capture population variability, or when transitioning towards biohybrid materials with cultured biological tissue, where large-scale replication is costly and resource intensive.

Researchers employing the tunable properties of organic hydrogels have developed a wide range of modeling approaches attempting to capture diffusion, deformation, and cross-linking behavior. These include continuum chemo-mechanical models, where multiphase frameworks couple reaction kinetics and swelling [33], finite-element simulations with detailed viscoelastic constitutive laws [34], and mesoscopic or kinetics-based models that link bond formation to macroscopic mechanics [29]. More recent statistical approaches apply Bayesian inference to soft materials, including hierarchical Bayesian model selection for constitutive behavior at high strain rates [35] and Bayesian optimization for nonlinear characterization of commercial and decellularized hydrogels [30]. While these methods provide valuable insight into local cross-linking dynamics, nonlinear deformation, and formulation-specific behavior, they often require extensive parameterization and narrowly defined gelation or loading conditions that limit their generality. Additionally, they typically assume steady-state mechanics and emphasize high-rate or transient deformation regimes. This focus contrasts with the low-strain, quasi-static deformation regimes often relevant for soft robotic actuation, as many aquatic and bioinspired systems operate primarily through low-strain bending or small-amplitude shape change rather than large bulk compression [3–6, 8–11]. Moreover, many of these approaches are not designed to accommodate batch-to-batch variability, fabrication-dependent effects, or population-level distributions that dominate functional performance in biologically-derived materials. For biodegradable or biohybrid robotic systems, these macroscopic behaviors, including steady-state stiffness, swelling, and variability across materials and fabrication workflows, must be characterized in a manner that is experimentally tractable, geometry-agnostic, and robust to inherent material heterogeneity.

To address these challenges, we introduce an accessible experimental–computational framework (Figure 1) designed to (1) explore polymer network variability in stock materials, (2) minimize variability during fabrication and cross-linking, (3) quantify the evolving material properties of printed hydrogels, and (4) capture sample-to-sample variability in the population. The workflow integrates molecular-level characterization of precursor composition via proton nuclear magnetic resonance (^1^H-NMR) spectroscopy, systematic measurement of swelling and stiffness across controlled cross-linking time and temperature conditions, and a generative model that characterizes and predicts population-level distributions and the expected range of material variability. Given the practical constraints of small sample sizes and single-timepoint material measurements, we calibrate this population model using a Synthetic-Likelihood Sequential Monte Carlo (SL-SMC) approach [36]. Although demonstrated here using embedded printing methods such as FRESH and using sodium alginate as a test case, the framework is fabrication-workflow- and material-agnostic, organized around population behavior rather than a specific formulation or technique. By providing a structured way to quantify variability at the molecular, fabrication, and bulk-mechanical levels, this framework enables more predictable use of biologically derived hydrogels in soft robotic actuators, tissue-mimetic constructs, and biodegradable devices.

**Figure 1.**
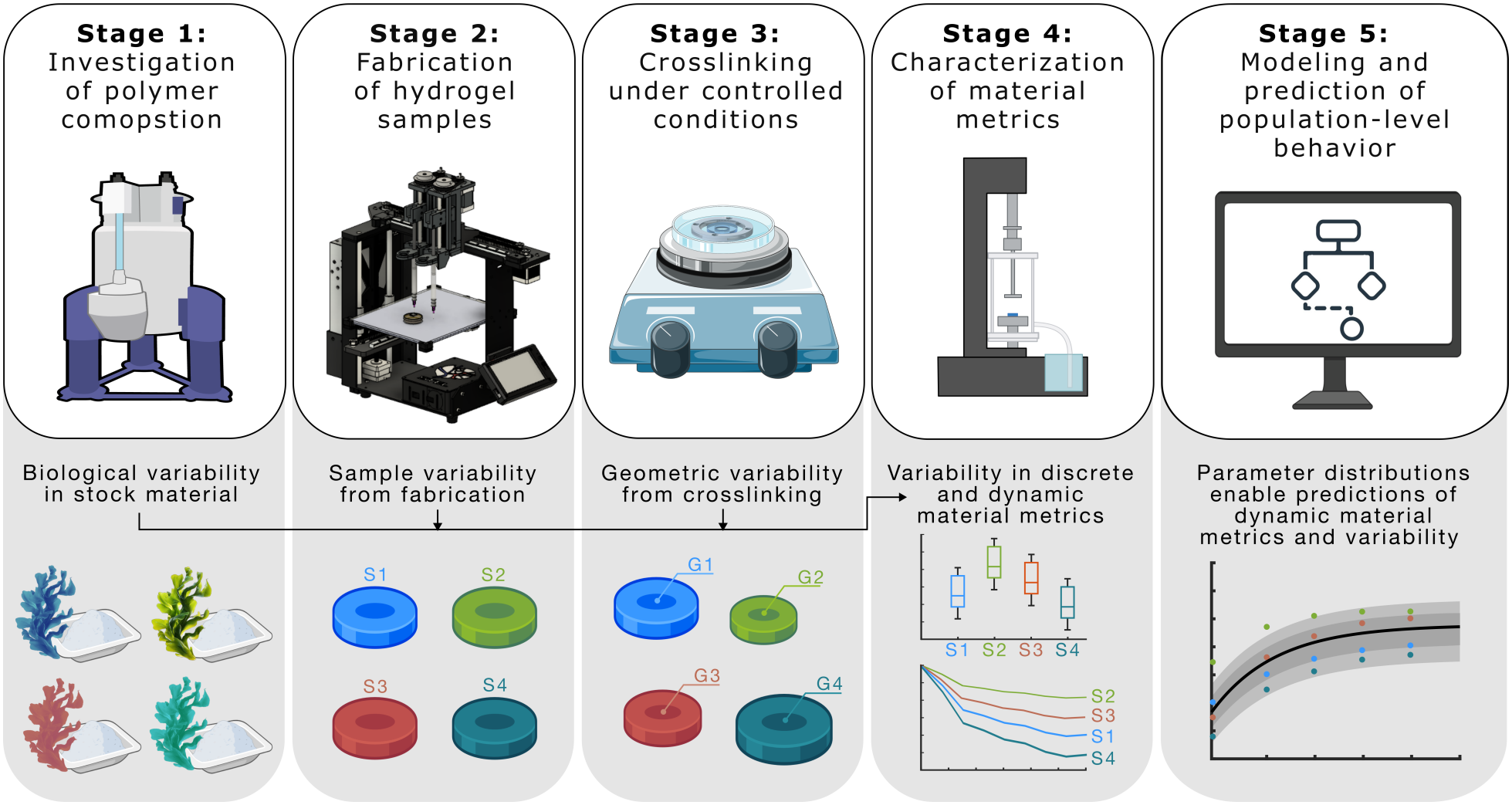
Experimental-computational framework to first investigate variability in material composition, fabrication methods, and cross-linking dynamics, characterize the impact on material metrics, and finally, model population-level distributions that quantify and predict variability. Bioprinter graphic courtesy of Shiwarski Tissue Engineering Lab Open Source Designs, and remaining equipment icons and graphics provided by Servier Medical Art (https://smart.servier.com), licensed under CC BY 4.0 (https://creativecommons.org/licenses/by/4.0/).

## Materials and Methods

### Investigation of Polymer Composition

#### ^1^H-NMR Spectroscopy of Stock Alginate Material

To prepare the alginate stock solution for ^1^H-NMR analysis (Figure 2), 0.5 g of sodium alginate powder (Allevi, A-0452) from 3 respective lots was added to a test tube of 12.5 mL of sterile deuterium oxide (D_2_O, heavy water) for a 4 % (w/v) solution. The tube was agitated in an orbital shaker for 1 hr until the powder was completely dissolved. 600 *µ*L of solution was transferred to a 5 mm ^1^H-NMR tube and loaded into a Bruker Systems NEO™ 500 MHz NMR Spectrometer for ^1^H-NMR spectroscopy at 333 K and 10 Hz. High-temperature spectroscopy was required to shift the dominating D_2_O spectral peak upfield, allowing the downfield region of monomer peaks to remain unoccluded. The resulting spectra were analyzed in Bruker TopSpin software.

**Figure 2.**
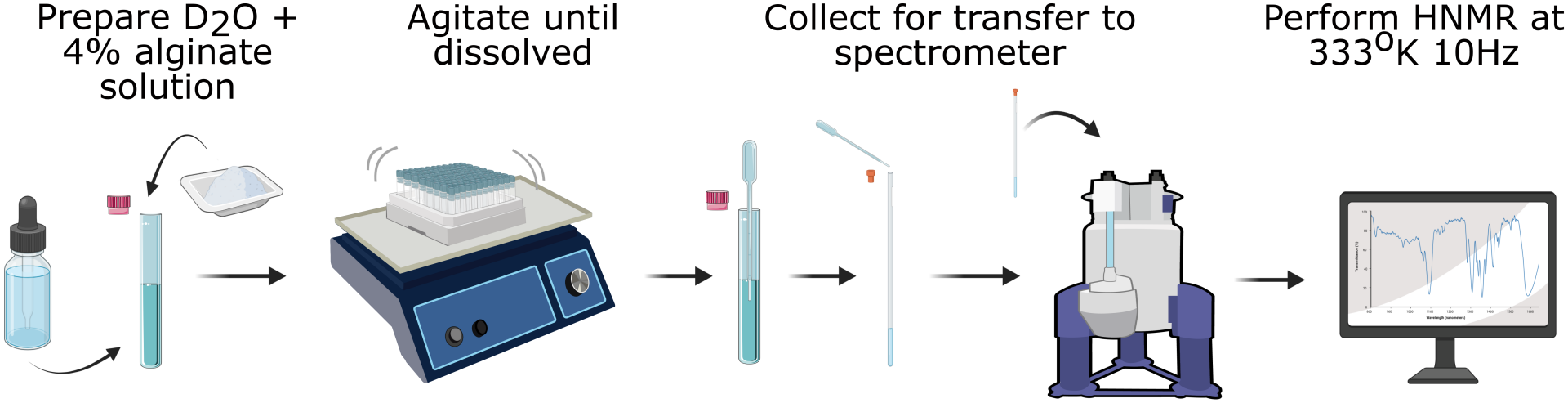
Preparation of sodium alginate ink for ^1^H-NMR spectroscopy. Following mixing of D_2_O and sodium alginate powder to achieve a 4 % (w/v) concentration, the solution was agitated, pipetted into a 5 mm ^1^H-NMR tube, and transferred to the spectrometer for high-temperature ^1^H-NMR. Equipment icons and graphics from NIAID NIH BioArt Source (bioart.niaid.nih.gov/bioart/).

#### Alginate Monomer Analysis

Methods for calculating monomer ratios were adapted from previous literature [26, 28, 37]. Briefly, peaks corresponding to Guluronic anomeric proton residues G1, Mannuronic acid dominated blocks (G[G5]M, M[G5]M, [M1]G, or [M1]M), and Guluronic acid dominated blocks (G[G5]G) were identified and assigned in the downfield spectra region. TopSpin’s internal deconvolution algorithm [38, 39] was used to disentangle overlapping peaks and approximate the integrals of each area. The integrals represent the frequency of G and M residues such that

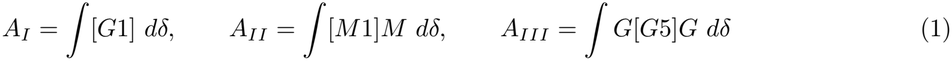

where *A*_*I*_ is the occurrence of G residues, *A*_*II*_ is the occurrence of M blocks, and *A*_*III*_ is the occurrence of G blocks. Given these assignments, the monomer fractions (*F*_*G*_ and *F*_*M*_) and resulting M/G ratio can be calculated given

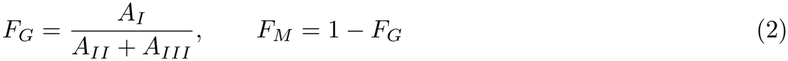

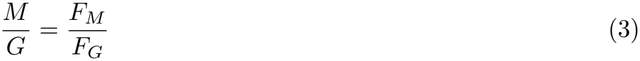

The M/G ratio was used as a quantifiable metric for the relative presence of each type of monomer in the alginate stock material and as an indication of its likelihood to cross-link fully in a predictable way. A lower ratio indicates a dominance of G blocks, which have a high affinity for calcium ions, and form predictable egg-box structures that translate to a stiffer and more uniform polymer network when ionically cross-linked [19, 26, 28, 37]. A higher ratio indicates a dominance of M blocks, whose irregular chain structures contribute to the elasticity of the network, but typically result in less uniform cross-linking. See Supporting Information; Equation S1 for calculation methods of additional metrics including monomer doublet fractions and ratios.

### Fabrication of Hydrogel Samples

#### Preparation of Sodium Alginate Ink for FRESH Printing

The alginate bioink was prepared according to previous works [11, 21, 22] by solubilizing sodium alginate powder (Allevi, A-0452) in heated (65°C) deionized water to a concentration of 4 % w/v and mixing using a magnetic stirrer. Alcian Blue powder (Alfa Aesar, J60122) was added to the solution before mixing at a concentration of 0.02 % (w/v) to increase visibility of the hydrogel during printing and imaging.

#### Preparation of Gelatin Support Bath for FRESH Printing

The gelatin support bath was prepared according to previous works [11, 21, 22] (Figure 3A). Briefly, 2.0% (w/v) Gelatin Type B (Fisher Chemical, G7-500), 0.25% (w/v) Pluronic F-127 (Sigma-Aldrich, F-127), and 1.0% (w/v) Gum Arabic (Sigma-Aldrich, G9752) were thoroughly mixed in 50% (v/v) ethanol/deionized water solution at 70–80°C using magnetic stirring. The solution was adjusted to pH 5.55 by adding 1N HCl dropwise while monitoring with a benchtop pH meter (Apera Instruments). This solution was stirred overnight in a temperature-controlled room (21°C) using an overhead stirrer at 550–575 rpm. After stirring, the gelatin slurry was washed three times with 0.1% (w/v) CaCl_2_ and stored at 4°C. Finally, the slurry was diluted to a concentration of 0.05 % (w/v) CaCl_2_, vortexed, and centrifuged at 2000 g for 5 min immediately before printing.

**Figure 3.**
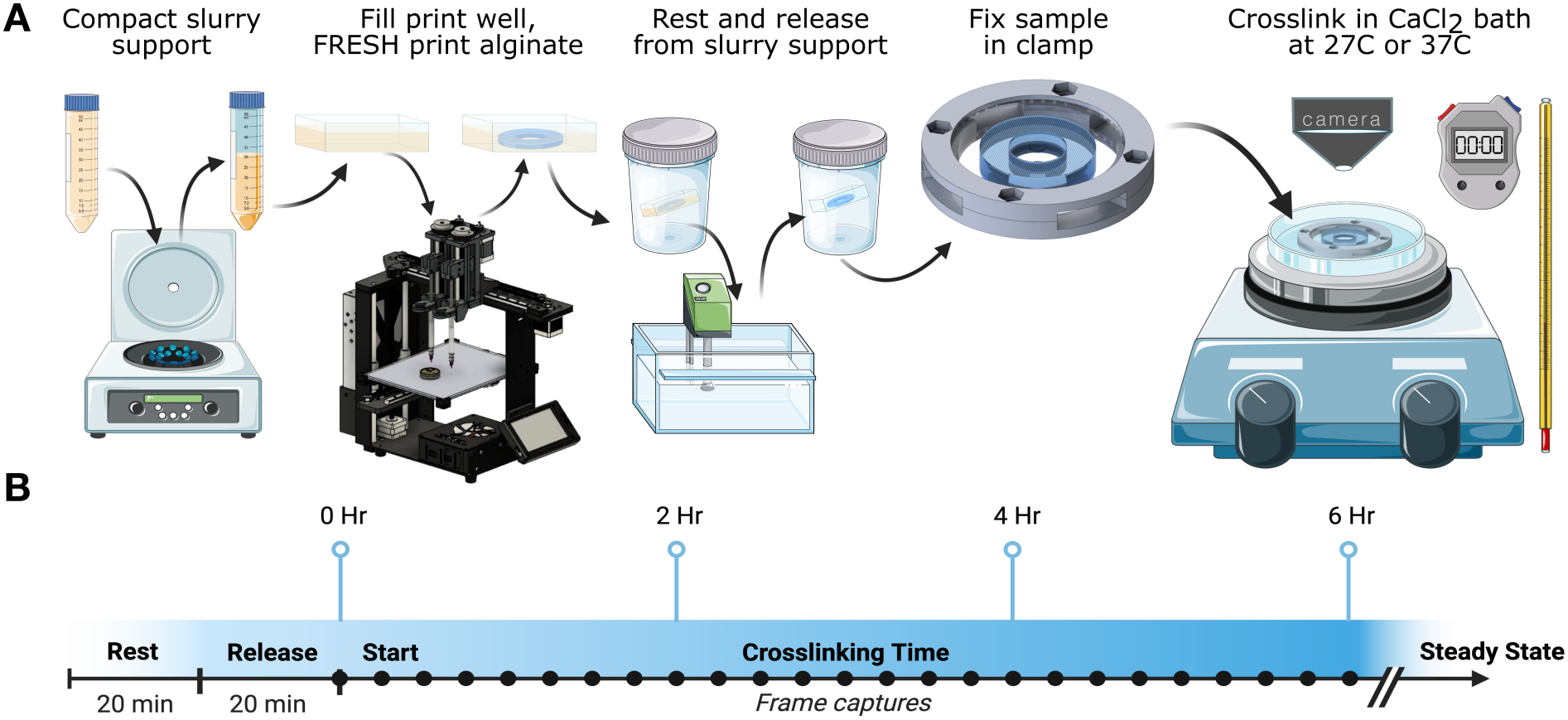
FRESH printing and cross-linking protocol. A) From left to right: Preparation of gelatin slurry for FRESH printing, FRESH printing of alginate geometry, post-print processing and slurry liquefaction, fixation of sample for cross-linking, cross-linking under controlled, monitored conditions. B) Timeline of post-printing processing and experimental cross-linking protocol. Bioprinter graphic courtesy of Shiwarski Tissue Engineering Lab Open Source Designs, and remaining equipment icons and graphics provided by Servier Medical Art (https://smart.servier.com), licensed under CC BY 4.0 (https://creativecommons.org/licenses/by/4.0/).

#### FRESH 3D Printing Protocol

Methods for the 3D printing of hydrogels were adapted from literature pioneering the FRESH method [21], with parameters refined for sodium alginate inks in soft robotic applications [11, 22] (Figure 3A). Annular rings with a 10.0 mm outer diameter, 5.0 mm inner radii, 5.0 mm height, cross-sectional surface area of approximately 235 mm^2^ and volume of approximately 1180 mm^3^ were printed for both swelling experiments and mechanical testing. All digital models were designed in SolidWorks (Academic License; Dassault Systémes) and converted to geometric code (G-code) using a customized configuration profile in Slic3r (http://slic3r.org) (20 mm/s print speed, 80 *µ*m layer height, 0.16 mm extrusion width, perimeters only). 3D printing was conducted with a Replistruder V4 [40] on a desktop Core XY 3D printer (Elf, Creativity Technology). To prepare for printing, the viscous sodium alginate ink was stored at 4°C and loaded into a 5 mL gas-tight syringe (Model 1005TLL, Hamilton) with a G30 blunt-tip needle (DN-05-LP-30, Bestean). Excess air bubbles were expelled mechanically by inverting and agitating the syringe. Printing wells were constructed from acrylic material to house the gelatin slurry suspension and fixed to the printer bed with double-sided tape.

During printing, alginate structures were partially cross-linked by the 0.05% (w/v) CaCl_2_ remaining in the slurry to ensure filament fusion. The FRESH printing time of the ring-shaped structure was 1 hr 30 min from the first to last layer. Immediately following printing, samples were incubated in the printing well at room temperature (27°C) for 20 min. The wells were then transferred to a dish with 2.5% (w/v) CaCl_2_ and incubated in a 37°C water bath for 20 min for slurry liquefaction. These processing steps are vital to the structural integrity and successful retrieval of FRESH-printed parts; in this experiment, they were all considered part of the pre-cross-link protocol (t < 0) (Figure 3B).

#### Cross-linking Protocol for Fabricated Samples

Following slurry liquefaction, baseline cross-linked samples were patted dry with a wipe (Kimtech Science™) to remove excess solution and weighed. The samples’ height was measured with calipers. Samples were then loaded into a 3D-printed (Prusa) PLA clamp such that the top and bottom of the sample were fixed between two acrylic plates (Figure 3A). The plates restricted perfusion through exposed surface areas to the inner and outer radial walls and restricted shape changes due to polymer network contraction to radial directions to facilitate longitudinal imaging.

To factorially investigate the impact of cross-linking time and temperature on alginate material properties, samples were cross-linked in two temperature groups (27°C or 37°C) in 4 discrete time groups (0 hr, 2 hr, 4 hr, or 6 hr) for a total of 7 test condition groups (N=6 samples per group; 0hr baseline cross-linked samples were not segregated by temperature). Each printed and clamped sample, after the required rest and release period, was submerged in a bath of 2.5 % CaCl_2_ cross-linking solution at t = 0 (Figure 3B). Bath temperature was controlled using a hot plate (Cole Parmer, 03404-39) and monitored with a temperature probe (Thermo Scientific, PT1000). Samples were monitored by a webcam (C920, Logitech) that recorded 1080p video and captured frames every 15 min. Initial and final diameters of the samples were measured from frames at *t* = 0 min and *t* = 120, 140 or 360 min for the 2 hour, 4 hour, and 6 hour cross-linking groups, respectively. Immediately after the assigned cross-linking time, samples were removed from the solution, patted dry with a wipe (Kimtech Science™), and reweighed.

### Characterization of Material Metrics

#### Swelling Measurement and Analysis

The impact of cross-linking time and temperature on sample swelling was quantified by tracking the change in the annular ring samples’ cross-sectional area over time. Because samples were restricted from motion in the height direction, all contraction was measured through inner and outer radii changes. Frames from the cross-linking videos were captured every 15 min and imported to MATLAB (MathWorks 2024b). Each frame was converted to an HSV pixel map, a mask was created to isolate the blue range of the dyed alginate, and a size threshold (< 10000 px) filter was applied to reduce noise (Figure 4A). The map was converted to a binary image for edge detection. Based on the average radii from the map center to the detected edges, circles were fit to the inner and outer edges of the annulus (Figure 4B), and the internal cross-sectional area of the sample was calculated.

**Figure 4.**
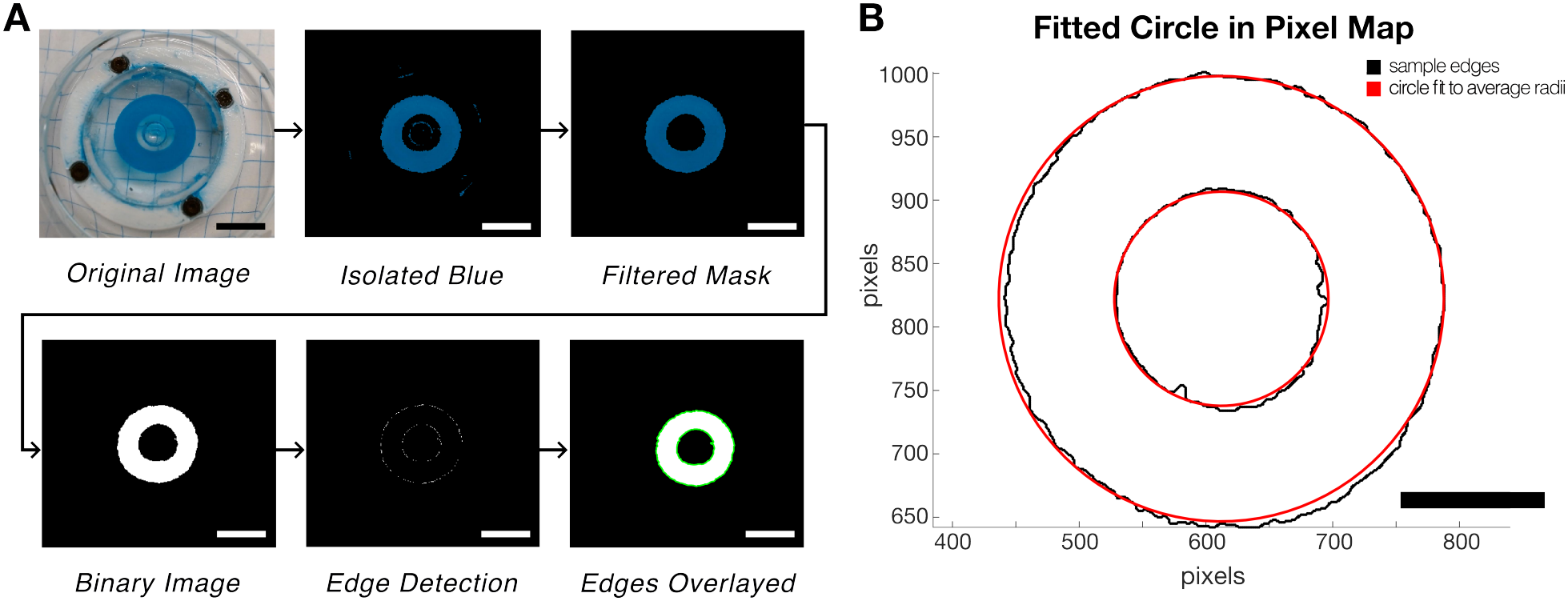
Methods for the characterization of material swelling during cross-linking. A) From left to right, top to bottom: Original image frame captured from camera feed, isolated HSV pixels in blue range, filtered mask to reduce noise, converted binary image, edges detected from binary transition boundary, edges overlaid with visual highlight. Scale bar: 10 mm. B) Highlighted circles of inner and outer circumference based on average radii distance from center to detected edge pixels. Scale bar: 5mm.

After N=6 samples had been tested for each cross-linking time and temperature group, the net change in mass after cross-linking was recorded as the percent mass reduction and analyzed according to cross-linking time and temperature.

#### Material Stiffness Testing and Analysis

Material stiffness was characterized using a universal testing system (MTS Systems, Criterion Model C41) with a biobath accessory equiped with a 50 N load cell (Figure 5A). The biobath was filled to a fixed height with 2.5% (w/v) CaCl_2_ solution for aquatic testing, and the crosshead was equipped with a compression plate fixture. An initial test cycle was performed without a sample present to measure ambient buoyancy forces in the biobath as the crosshead moved; this allowed us to fit a linear bouyancy force vs. displacement calibration curve. cross-linked alginate samples were fixed to the bottom of the testing stage using double-sided tape and water-bonding superglue (Loctite). The crosshead plate was lowered to be in contact with the top of the sample until a pre-load force of 0.10 mN was detected. Samples were then cyclically compressed 1.5 mm at a velocity rate of 0.1 mm/s for 30 cycles until the peak forces measured converged (Δ*F*_*peak*_ *≤* 5%) (Figure 5B). Force and crosshead displacement data were recorded at 10 Hz and exported for analysis in MATLAB (MathWorks 2024b).

**Figure 5.**
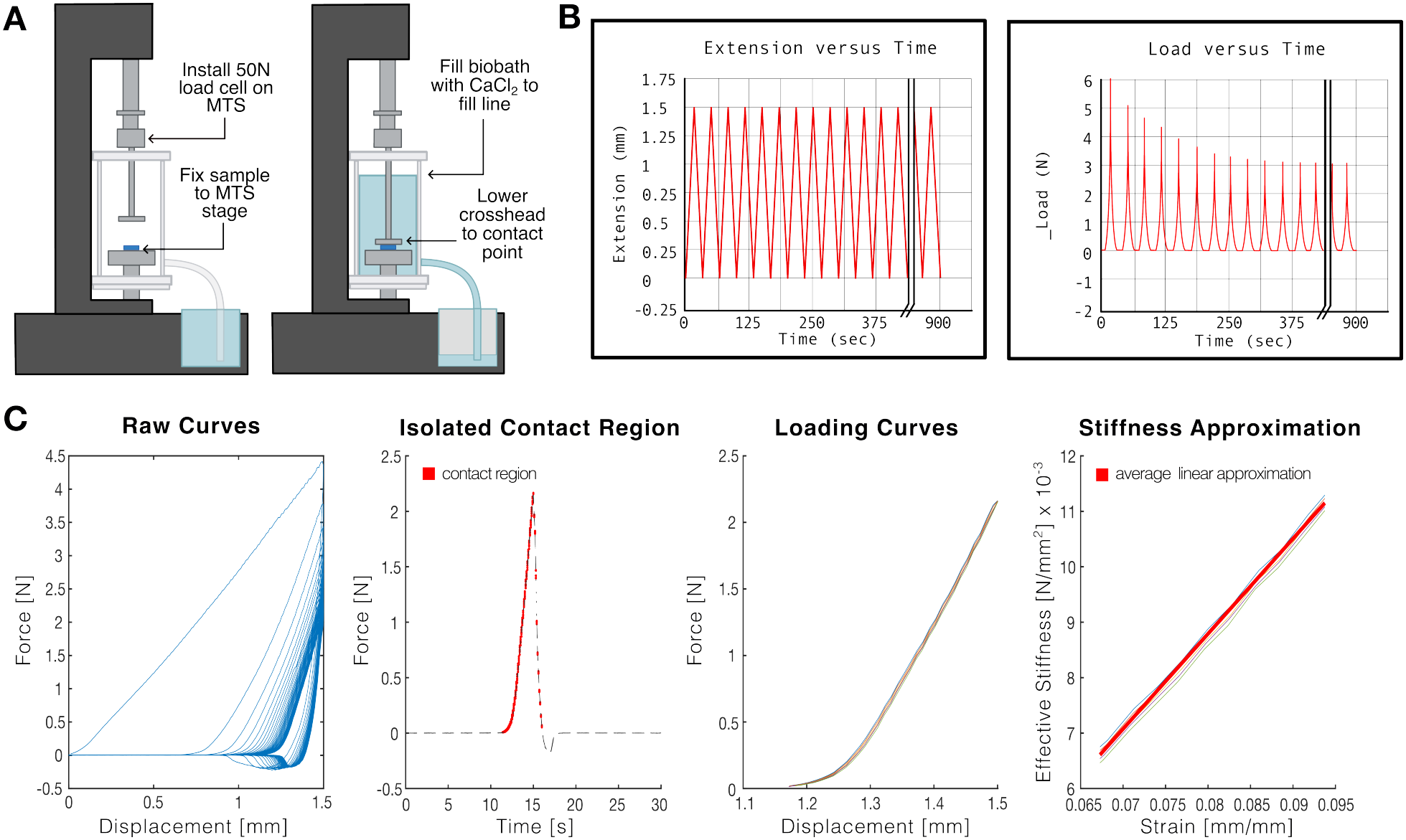
Methods for the characterization of cross-linked material stiffness. A) Fixation of printed alginate sample on the stage of a material testing system (MTS) equipped with a 50N load cell. The biobath is filled with CaCl_2_ to a fixed height before lowering the crosshead into contact position. B) Extension and Load curves of the programmed MTS cyclic compression for 30 cycles at a strain rate of 0.1 mm/s with a 10z sampling frequency. C) From left to right: Raw data force vs. displacement curves of a representative sample (2 hr @ 27^*°*^C, sampleID: S2c) after buoyancy and noise floor correction, isolated contact regions of the loading curve after buoyancy and noise correction, loading curves of last 5 cycles after convergence Δ *≤* 5%, average secant linear approximation of 0.07 *≤ ε ≤* 0.10 strain region.

In MATLAB, the force vs. displacement data was initially corrected for ambient noise and buoyancy forces; buoyancy forces were subtracted at each displacement level using the linear calibration curve, and any non-mean-zero noise floor was subtracted from the data to reset the zero-load condition. Next, the loading curves of the last 5 cycles after convergence were isolated (Figure 5C). Plastic deformation in the material from pre-straining in the initial cycles resulted in a region of the curve where the crosshead was not in contact with the sample. This zero-contact region spanned approximately 1.1 mm in the converged cycles. Therefore, curves were truncated to only include contact regions, and sample dimensions were corrected to account for plastic deformation before stress *σ* (N/mm^2^) and strain *ε* (mm/mm) calculations. Specifically, the effective rest length *L*_0_ was reduced by the measured deformation offset (Δ*L*_*plastic*_, and the cross-sectional area was scaled by the corresponding stretch ratio (*A* = *A*_0_*/λ*_0_, where *λ*_0_ = 1 *−* Δ*L*_*plastic*_*/H* and *H* is the rest height of the sample prior to testing). The resulting stress region occurred over a 10% strain region in the plastically deformed material. An end-of-cycle (0.07 *≤ ε ≤* 0.10) secant linear approximation of the converged stress-strain slopes was calculated using the mean stress-strain curve fit to the last 5 cycles (Figure 5C). This mean slope represents an effective elastic modulus *E*_*eff*_ of the viscoelastic material to be used as a comparative metric between samples under various controlled cross-linking conditions.

The characterization of stiffness using this compression-based metric relies on simplifying assumptions, including Hookean behavior, incompressibility, and isotropy. These assumptions do not capture the transient, rate-dependent, or nonlinear mechanics of ionically cross-linked hydrogels, nor do they represent a full constitutive description of the material. Rather than attempting to fit a detailed viscoelastic model, this quasistatic approximation provides a consistent and tractable way to compare samples and quantify how cross-linking conditions influence average stiffness in the low-strain regime relevant to macroscopic robotic actuation. We therefore do not interpret *E*_*eff*_ as an absolute material property or a substitute for a full mechanical model; instead, it serves as a simplified, reproducible metric within the broader workflow to assess relative differences in steady-state behavior and material variability across batches and fabrication conditions. If complete constitutive models were required for a given application, additional mechanical experiments could be incorporated into this pipeline.

After N=6 samples had been tested for each cross-linking time and temperature group, the effective moduli were plotted according to cross-linking time and temperature.

### Population Modeling of Hydrogel Material Properties

Many hydrogel properties evolve over time as the material undergoes physical or chemical processes including swelling, contraction, solvent exchange, degradation, and cross-linking [19, 23–25, 27]. As a first-order approximation, we assume that the evolution of a measured property can be represented by first-order dynamics approaching a steady-state value. Let *y*_*i*_(*t*) denote the value of a property measured from sample *i* at time *t*. The temporal evolution of the property is modeled as

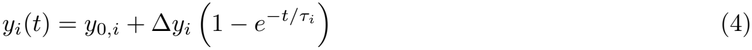

where *y*_0,*i*_ is the initial value of the property, Δ*y*_*i*_ is the total change in the property during the process of interest, and *τ*_*i*_ is the characteristic time constant governing the dynamics. Depending on the application, *y*_*i*_(*t*) may represent geometric, mechanical, chemical, or biological quantities. Additionally, while we assume first-order dynamics for our material, if the underlying mechanisms driving property changes are known, a more specific dynamical model could be utilized within this same framework.

The final properties of our hydrogel in this work may be affected by both the cross-linking duration and temperature, and the natural variability inherent in biologically derived materials (See ^1^H-NMR characterization reveals intrinsic variability in alginate monomer composition). To capture how material metrics evolve and account for specimen-to-specimen variability, we developed a generative statistical model in which the dynamic parameters describing each sample (*y*_0,*i*_, Δ*y*_*i*_,, and *τ*_*i*_) are treated as random variables drawn from an underlying population distribution. In this population model, *y*_*i*,0_ and Δ*y*_*i*_ are sampled from a joint Normal distribution

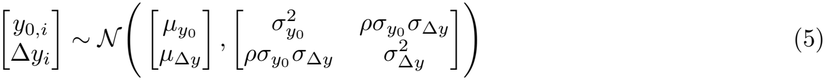

where *µ*_*J*_ and *σ*_*J*_ denote the population mean and standard deviation of each parameter *J*, and *ρ* represents the correlation between the initial value and total change. A joint Normal distribution (rather than two independent Normal distributions) was chosen to allow for coupling between the starting material state and the change in material properties caused by, e.g., the monomer ratio in the material. Finally, we assumed that the time constant of the cross-linking dynamics is independent of the stiffness and distributed as a Gamma random variable

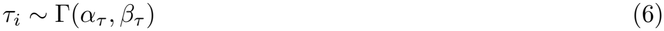

where *α*_*τ*_ and *β*_*τ*_ are the shape and scale parameters of the Gamma distribution. This framework could be extended to include coupling between the starting/change parameters and the time constant, but this is beyond the scope of this work. For easier interpretation, we parameterize the Gamma distribution by its mean *µ*_*τ*_ and standard deviation *σ*_*τ*_, such that *α*_*τ*_ = (*µ*_*τ*_ */σ*_*τ*_)^2^ and 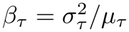. A Gamma distribution was chosen for the time constant to ensure that the time constant was positive. The full population model for the dynamics parameters governing the time course of a given material property *y*_*i*_(*t*) is then

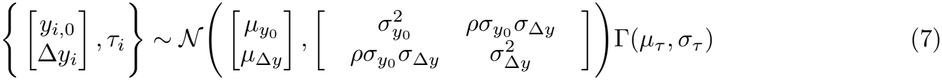

The parameters of this population model are captured in the parameter vector 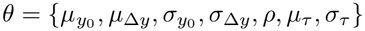.

A key assumption of this framework is that the majority of variability observed in hydrogel properties comes from sample-to-sample variability rather than from noisy measurements of those properties. For the hydrogel samples and measurement methods used in this study, this is a reasonable assumption. If low-noise measurements of hydrogel properties cannot be reliably obtained, this generative model could be coupled with an additional noise model, for which parameters would also need to be estimated. This noise model is not considered here.

#### Conditional Effect Parameters to Account for Temperature Conditions

To account for post-processing-dependent behavior, we extended the population model with conditional effect parameters for the change term, Δ*y*, and the characteristic time constant, *τ*. For our experiments here, that post-processing condition is the cross-linking temperature, but a similar approach could be used to estimate the effects of other conditions, such as cross-linker chemistry or concentration. Additionally, the parameters associated with the starting condition (*µ*_*y*0_ and *σ*_*y*0_) are not considered, as they are estimated from data collected before extended cross-linking begins and are not influenced by cross-linking conditions.

Separate effect parameters were included for the means and spreads of both Δ*y* and *τ*, allowing temperature to independently influence the expected magnitude and variability of the observed response. This approach preserves a common model structure across conditions while permitting post-processing-specific changes in both the dynamics and heterogeneity of the population.

The temperature-conditioned population parameters are defined as

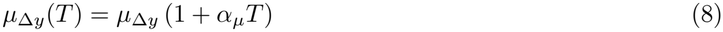

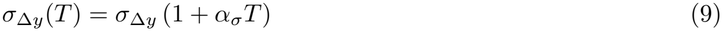

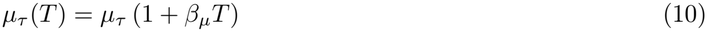

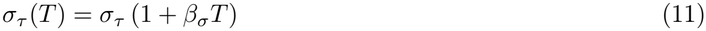

where *α*_*µ*_ and *α*_*σ*_ describe temperature-dependent changes in the mean and standard deviation of Δ*y*, respectively, and *β*_*µ*_ and *β*_*σ*_ describe temperature-dependent changes in the mean and standard deviation of *τ*. The indicator variable *T* takes a value of 0 for the lower-temperature condition and 1 for the elevated-temperature condition, such that the baseline parameters correspond directly to the lower-temperature population. However, this same framework could be extended to account for continuous variables as well if several conditions were tested. Substituting these terms into the population model yields

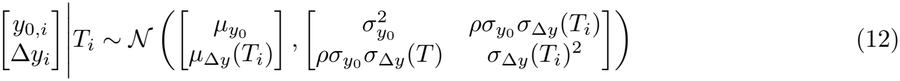

and

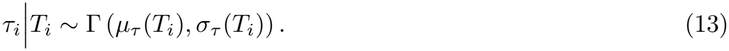

The expanded population-level parameter vector describing material properties is now

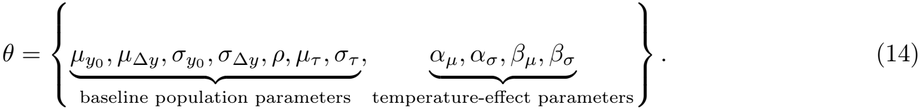

#### Calibration of Population Model Parameters using Sequential Monte Carlo Estimation

For this statistical model, we aim to estimate the population model parameter vector from observed properties data. Here, we chose to utilize a Bayesian framework for this calibration. Therefore, we aim to estimate the posterior distribution of the population model parameters (*θ*) given the observed data ({*y*_*i*_(*t*_*j*_)}) using Bayes’ Theorem:

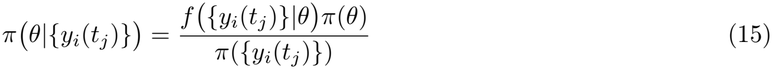

where *π (θ*|*{y*_*i*_(*t*_*j*_)}) is the posterior distribution of interest, *f({y*_*i*_(*t*_*j*_)} | *θ*) is the likelihood based on our population model, *π*(*θ*) is the prior of our model parameters, and *π*({*y*_*i*_(*t*_*j*_)}) is the total probability of the dataset, which serves as a normalizing constant. We chose to utilize a Bayesian framework for this framework (as opposed to a Frequentist approach utilizing Maximum Likelihood Estimation) because it 1) allows us to bring in information from previous experiments to inform our parameter estimation through our choice of priors, 2) is naturally set up for estimation refinement through new experiments, and 3) directly gives us measures of parameter uncertainty. Additionally, a Bayesian approach provides a suite of computational tools for estimating the model parameters that are amenable to the various types of datasets that we will collect during these experiments. Specifically, for this framework, we utilized Sequential Monte Carlo (SMC) methods [41–43], in which a set of particles (*{θ*^*\**^*}*) is used to estimate the posterior progressively by passing them through a series of intermediate distributions. Through this series of distributions, more probable particles are more heavily weighted, and less probable ones are weighted less. For details of the SMC algorithms used in this framework, see the Supporting Information.

We cannot directly calculate the likelihood of our observed data (*f {y*_*i*_(*t*_*j*_)} | *θ*), as the population model only deals with the parameters of the exponential model (Eqn. 4). For datasets containing sufficient repeated measurements over time, such as the dynamic cross-sectional area (CSA) change, we can directly estimate those exponential model parameters and thus create a parameter dataset {*y*_0,*i*_, Δ*y*_*i*_, *τ*_*i*_} from the observed dataset {*y*_*i*_(*t*_*j*_)}. For the deterministic exponential model, we can use standard curve fitting tools to estimate these parameter datasets (here, nonlinear total least squares regression using MATLAB’s *fminsearch* function). Then, the likelihood can be calculated for this parameter dataset, and standard SMC methods can be used (see Supporting Information: Summary of Sequential Monte Carlo Methods and Implementation).

However, some hydrogel properties can not be monitored continuously because their measurement requires fully or semi-destructive testing. Examples include effective elastic modulus and mass reduction measurements, where each specimen contributes only a single observation at a particular time point. Under these conditions, we do not have independent information for *y*_0,*i*_, Δ*y*_*i*_, and *τ*_*i*_ from each sample, and therefore, we cannot calculate our parameter dataset. Accounting for this incomplete information directly in the calculation of the likelihood would require integration over a high-dimensional space and quickly becomes difficult. To extend the population modeling framework to such datasets, the SMC algorithm was combined with Synthetic Likelihood methods (SL-SMC). In SL-SMC, as in other Approximate Bayesian Computation (ABC) methods [36, 42, 44], the likelihood of a dataset for a given model cannot be easily calculated, but synthetic data from that model can easily be sampled. Using these synthetic data, we can infer how likely our observed data are based on how similar the two datasets are to each other. In ABC methods, this similarity is captured through a distance function, and the intermediate distributions of the SMC sampler progressively shrink the error tolerance of that distance function. In SL methods, the distribution of the data or of summary metrics of the data is directly estimated using the synthetic data. While this can be achieved in a variety of ways, here we utilized Gaussian kernel density estimation [45, 46], where the probability of the data is estimated as the sum of many Normal distributions centered on our synthetic data. Thus, from the synthetic data 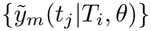, the probability of a particular value *y*_*i*_(*t*_*j*_|*T*_*i*_) is calculated as

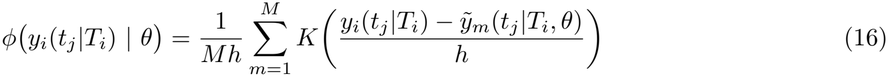

where *M* is the number of synthetic data points that were generated, *K*(*·*) is the standard Normal distribution probability density function, and *h* is the bandwidth of the density estimate. Here, we set the bandwidth using Silverman’s rule of thumb [47]. Then, the synthetic likelihood estimation becomes

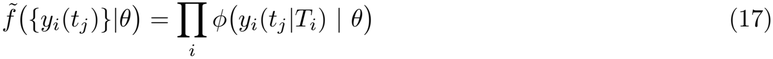

A major benefit of SL methods over other ABC methods is that exactly the same SMC algorithm can be used, but with the synthetic likelihood 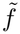 replacing the true likelihood *f*. In ABC methods, the weight updating rule and resampling strategies must be modified.

The SMC and SL-SMC methods provide the calibrated joint posterior for the population model parameters and allow for inference on those parameters. For all parameters, we report the posterior median and 95% credible interval (the region over which there is a 95% chance of finding the underlying parameter). Using the posterior median model parameters, we can also construct the posterior median population model and simulate dynamic trajectories of our property of interest. To calculate all posterior dynamic trajectory summaries (mean, standard deviation, and 95% prediction interval of the trajectory distribution at the posterior median parameters), 100,000 simulations were conducted.

Detailed methods for the implementation of the SMC and SL-SMC sampler for each experimental material metric can be found in the Supporting Information. This includes both the algorithms used by the samplers and the definitions of the priors used for each dataset. A MATLAB library for SMC sampling (using either true or synthetic likelihoods), along with validation scripts, was created and is available on GitHub (See data availability). In addition, a class implementation for our population model is similarly available. Finally, the GitHub contains all code needed to perform the parameter dataset calculation, the posterior calibration, and posterior inference for each dataset. All code was written in MATLAB R2024b, and all parameter estimation and simulations were conducted using an AMD Ryzen 7 processor (3.30 GHz, 8-cores, 16GB RAM) running Windows 11. The full calibration across all datasets (including posterior predictive simulation) can be completed in approximately 1 hour (8 core hours).

## Results and Discussion

In this work, we develop a combined experimental and computational framework for characterizing how material properties of 3D-printed hydrogels change during crosslinking and how these dynamics vary within a population. This tool allows for the quantitative investigation of bio-derived material crosslinking and material variability across spatial and temporal scales. Using a population statistical model in combination with a first-order dynamics model, we can quantify the variability in steady-state and dynamical material properties and predict the variability in future parts. Here, we present the application of the framework to FRESH-printed sodium alginate constructs as a case study. Results are presented in a progression from material characterization to population-level modeling. First, ^1^H-NMR analysis quantifies variability in the stock alginate material prior to fabrication. Next, geometric, mechanical, and mass-transfer measurements from cross-linking experiments establish the observed sample-to-sample variability in hydrogel behavior. We then incorporate these data into the computational modeling framework to estimate population-level distributions, dynamic responses, and parameter uncertainty. Finally, we compare trends across all modeled properties to provide insight into hydrogel cross-linking behavior and illustrate the value of the combined experimental-computational workflow.

### ^1^H-NMR characterization reveals intrinsic variability in alginate monomer composition

H-NMR spectroscopy of three alginate samples, followed by spectral deconvolution, yielded clear and consistent peak assignments for the characteristic monomer residues (G and M) and their corresponding block structures (Figure 6A and B). These assignments enabled direct calculation of the M/G monomer ratio and the MM/GG doublet fractions (Table 1), providing a molecular-level view of the intrinsic variability present in the stock materials.

**Table 1:** H-NMR peak areas and monomer ratios for alginate samples.

|  | **A<sub>I</sub>** (×10<sup>10</sup>) | **A<sub>II</sub>** (×10<sup>10</sup>) | **A<sub>III</sub>** (×10<sup>10</sup>) | $\frac{\mathbf{M}}{\mathbf{G}}$ | $\frac{\mathbf{MM}}{\mathbf{GG}}$ |
|---|---|---|---|---|---|
|  | (*a.u.*) | (*a.u.*) | (*a.u.*) |  |  |
| Sample 1 | 39.32 | 54.51 | 6.549 | 0.5530 | 1.6838 |
| Sample 2 | 62.44 | 74.23 | 25.89 | 0.6034 | 0.0437 |
| Sample 3 | 51.88 | 72.21 | 26.61 | 0.9048 | 0.8144 |

**Figure 6.**
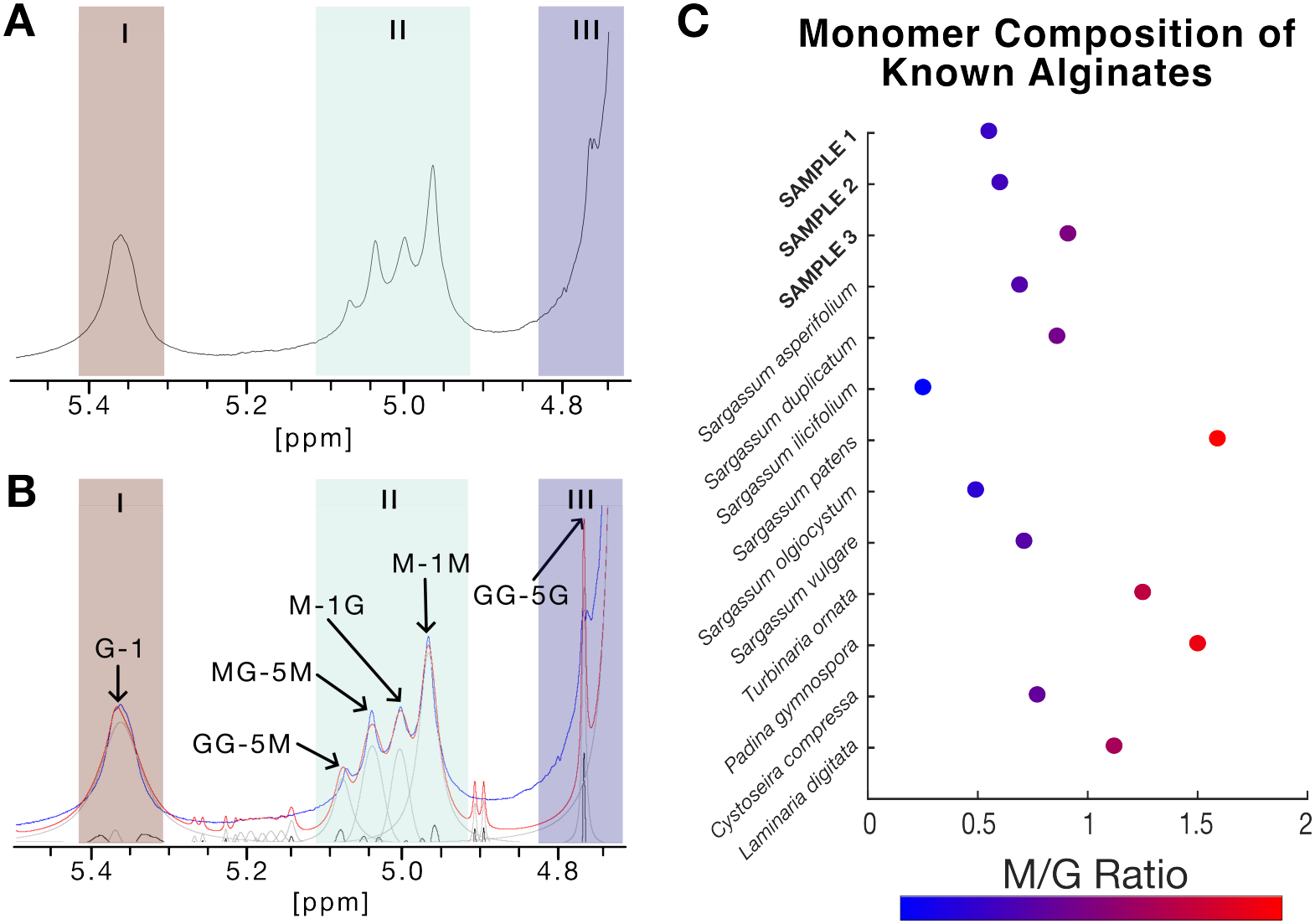
Results of alginate ^1^H-NMR spectral analysis. A) Representative sample (Sample 1) downfield region of raw spectra from Bruker’s Topspin software, with *D*_2_*O* peak appearing at approximately 4.7 ppm (off right) and monomer spectral region appearing from approximately 5.4 ppm to 4.75 ppm. B) Result of Topspin deconvolution algorithm for the same representative sample as shown in A (see full sample spectra and ratios in Supporting Information; Figure S1, Tables S1, S2), with assigned peaks corresponding to (I) Guluronic anomeric proton residues G1, (II) Mannuronic acid blocks [M1]M, and (III) Guluronic acid blocks G[G5]G. C) M/G ratio of tested samples (N = 3) compared to known ratios of seaweed species commonly harvested for the formulation of alginate stock material [26]. Note: both the horizontal axis in C) and the color scale represent the same M/G ratio data.

The measured M/G ratios for all three samples fell on the lower end of the M/G ratio range (< 1) reported for naturally sourced alginates [26] (Figure 6C), indicating a higher relative abundance of G blocks. Because higher G-block content increases cross-link density and contraction, while M-rich compositions yield softer, more extensible gels with slower gelation rates, these lower M/G ratios are broadly consistent with the stiffer material properties desirable in cross-linkable hydrogel fabrication [19, 27]. However, the sample-to-sample variance in M/G ratio was substantial, suggesting that we should expect to see a similarly substantial variance in the experimentally measured stiffness distributions of our printed samples.

More pronounced variability emerged in the MM/GG doublet fractions, which quantify the frequency of repeating M–M and G–G blocks. These doublet ratios varied dramatically across samples (spanning two orders of magnitude), indicating that the block repetition within the polymer, not just the overall monomer fraction, differs considerably between alginate batches. Because block structure governs the density, uniformity, and cooperativity of ionic cross-linking [19, 26–28], these differences likely propagate upward into the heterogeneous contraction and viscoelastic behavior observed across identically fabricated hydrogels.

Together, these results demonstrate that alginate’s molecular composition is inconsistent, even within a single supplier or chemical lot. This intrinsic variability poses a fundamental challenge for fabricating reproducible hydrogel inks for soft actuators.

### Effect of cross-linking conditions on cross-sectional area

#### Dynamic raw data reveal uncertainty in population-level behavior

To evaluate how cross-linking conditions influence geometric evolution, the cross-sectional area of each sample was tracked throughout the cross-linking period. Summaries of the trajectories normalized by their initial value for all experimental groups are shown in Figure 7. Despite substantial variability between samples, the measured responses consistently exhibited behavior characteristic of first-order dynamics, with the normalized cross-sectional area decaying exponentially over time toward a non-zero steady-state value.

**Figure 7.**
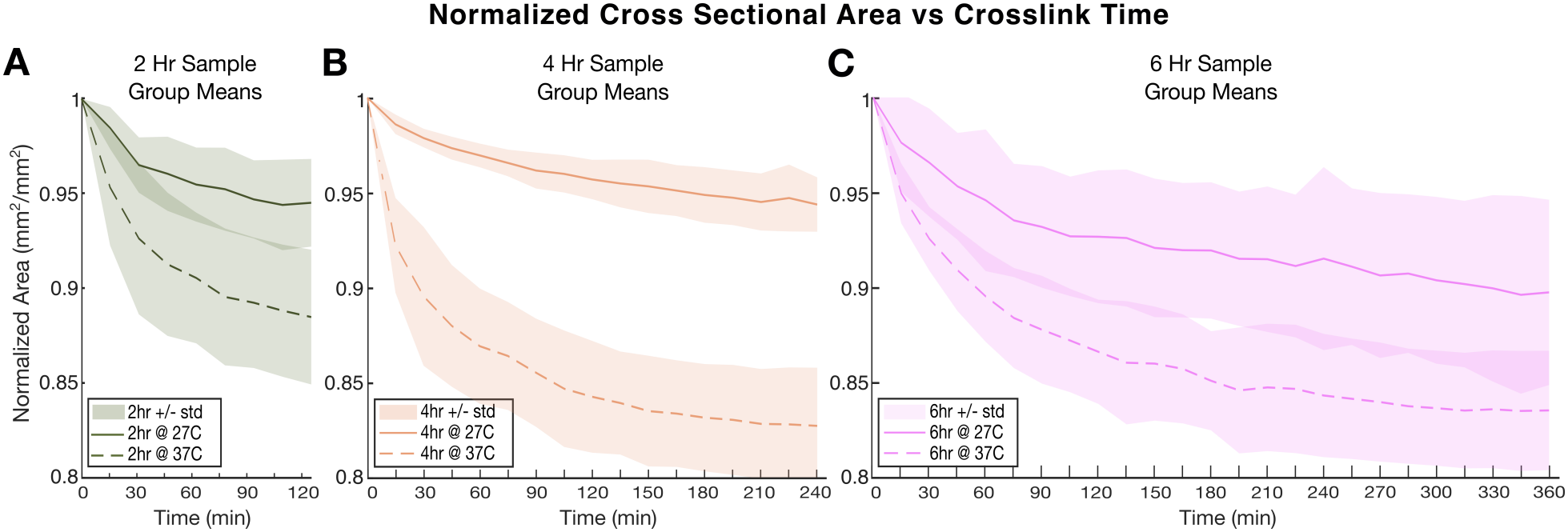
Effects of controlled cross-linking conditions on sample geometry. A) Normalized cross-sectional area of samples over the cross-linking period for both temperature conditions within the 2 hr group. B) Normalized cross-sectional area of samples over the cross-linking period for both temperature conditions within the 4 hr group. C) Normalized cross-sectional area of samples over the cross-linking period for both temperature conditions within the 6 hr group. Trendlines and shading represent mean and std, respectively, of all samples within a test group (N=6). Solid and dashed trendlines represent sample groups cross-linked at 27^*°*^C and 37^*°*^C, respectively.

To verify this observation, each individual sample was independently fit to the first-order constitutive model introduced in Eqn. 4, specialized here for normalized cross-sectional area,

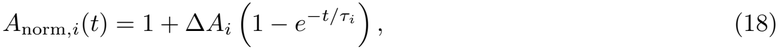

where *A*_norm,*i*_ is the normalized cross-sectional area of sample *i*, Δ*A*_*i*_ represents the total fractional change in area, and *τ*_*i*_ is the characteristic time constant governing the contraction dynamics. Individual sample fits are shown in Figure S2, and the corresponding fitting statistics are reported in Table S3. Across all experimental samples, the first-order model provided an accurate description of the observed individual sample dynamics, with consistently high coefficients of determination (average across samples in a group 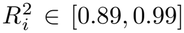. In contrast, fitting a single exponential model to all samples within an experimental group (N=6) resulted in substantially poorer agreement with the measured trajectories. Although the average response retained a generally exponential shape, the spread in contraction magnitude and contraction rate between samples produced large residual errors 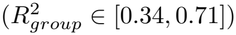 that could not be captured by a single deterministic fit.

This observation, that individual sample models fit well but that group level models are less informative, indicates that uncertainty in the measured response is dominated by population-level variability rather than by inadequacies in the constitutive model itself. Additionally, substantial overlap was observed between the distributions of normalized area trajectories measured under different temperature conditions (Fig. S2). Conventional Frequentist statistical analysis methods, which typically concentrate on group means and treat variability as noise rather than as a feature of the distribution, would provide limited insight into whether temperature impacts cross-linking without substantial sample sizes. More importantly, such methods do not explicitly characterize the broad variability observed within each experimental group, which is needed for forward prediction of material properties. This population-level variability and weakness of standard Frequentist methods motivated the use of a Bayesian population-based modeling framework capable of explicitly accounting for biological variability.

#### Population modeling of cross-sectional area enables prediction of population-level geometric fidelity and variability

To address the need to characterize population-level variability between samples, we developed a population statistical model capturing the dynamical parameters of our first-order model of the cross-sectional dynamics. To avoid the shortcomings of Frequentist analysis, we calibrated the parameters using a Bayesian framework and calculated the parameter posteriors using a developed Sequential Monte Carlo toolbox (Comparisons of marginal priors and posteriors as a result of the Sequential Monte Carlo inference process are shown in Figure 8A. Joint posteriors, probability density contours, and convergence over SMC generations are provided in the Supporting Information; Figure S4.). The posterior distributions (Table 2) reveal several important features of the underlying geometric variability within the hydrogel population. First, substantial variability was present in the baseline construct geometry, as indicated by the relatively large value of 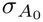 (29.74 [23.71, 38.69] mm^2^) relative to the mean initial cross-sectional area, 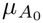 (198.87 [188.72, 208.87] mm^2^). This result is consistent with the broad distributions observed experimentally in Figure 9C. It is worth noting that because low-concentration ambient ionic diffusion is vital to the printing process, and because samples must be released from slurry into a high-concentration ionic solution before physical analysis is possible, the baseline construct geometry is not solely representative of the printing fidelity. Rather, samples have already undergone some degree of cross-linking at *t* = 0, making variability and shape deviation due to the FRESH manufacturing process indistinguishable from biological variability affecting polymer network contraction at this stage. In addition, some samples exhibited larger than designed diameters at the time of measurement despite later undergoing the expected contraction. This is likely an artifact of the viscous ink spreading and swelling after extrusion, the extent of which is dependent on manufacturing conditions such as ink temperature and slurry particle size, as well as the biological variability of the polymer network that affects ink viscosity and rheology. This modeling framework could be applied to alternative fabrication methods or to the cross-linking of materials that undergo less bulk shape change during polymer network formation, to better isolate the effects of the material variability and characterize the printing fidelity of the manufacturing process method itself.

**Table 2:**
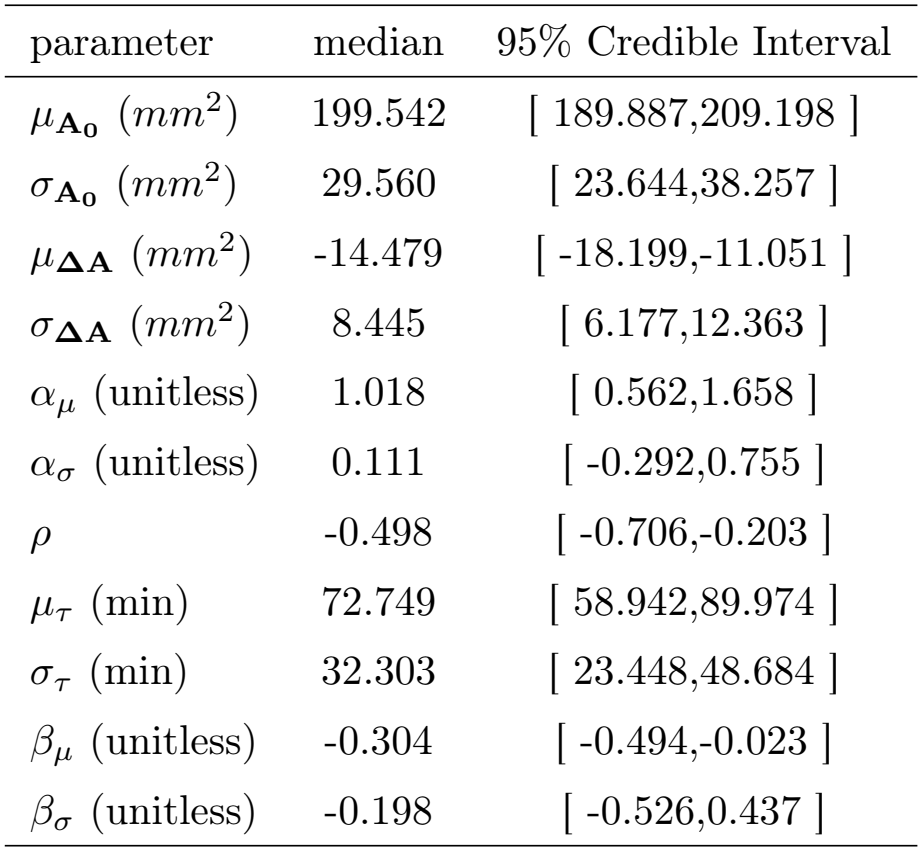
Posterior parameter outputs from converged Cross-sectional area population model.

| parameter | median | 95% Credible Interval |
|---|---|---|
| μ<sub>**A<sub>0</sub>**</sub> (*mm*<sup>2</sup>) | 199.542 | [ 189.887,209.198 ] |
| σ<sub>**A<sub>0</sub>**</sub> (*mm*<sup>2</sup>) | 29.560 | [ 23.644,38.257 ] |
| μ<sub>**ΔA**</sub> (*mm*<sup>2</sup>) | -14.479 | [ -18.199,-11.051 ] |
| σ<sub>**ΔA**</sub> (*mm*<sup>2</sup>) | 8.445 | [ 6.177,12.363 ] |
| α<sub>μ</sub> (unitless) | 1.018 | [ 0.562,1.658 ] |
| α<sub>σ</sub> (unitless) | 0.111 | [ -0.292,0.755 ] |
| ρ | -0.498 | [ -0.706,-0.203 ] |
| μ<sub>τ</sub> (min) | 72.749 | [ 58.942,89.974 ] |
| σ<sub>τ</sub> (min) | 32.303 | [ 23.448,48.684 ] |
| β<sub>μ</sub> (unitless) | -0.304 | [ -0.494,-0.023 ] |
| β<sub>σ</sub> (unitless) | -0.198 | [ -0.526,0.437 ] |

**Figure 8.**
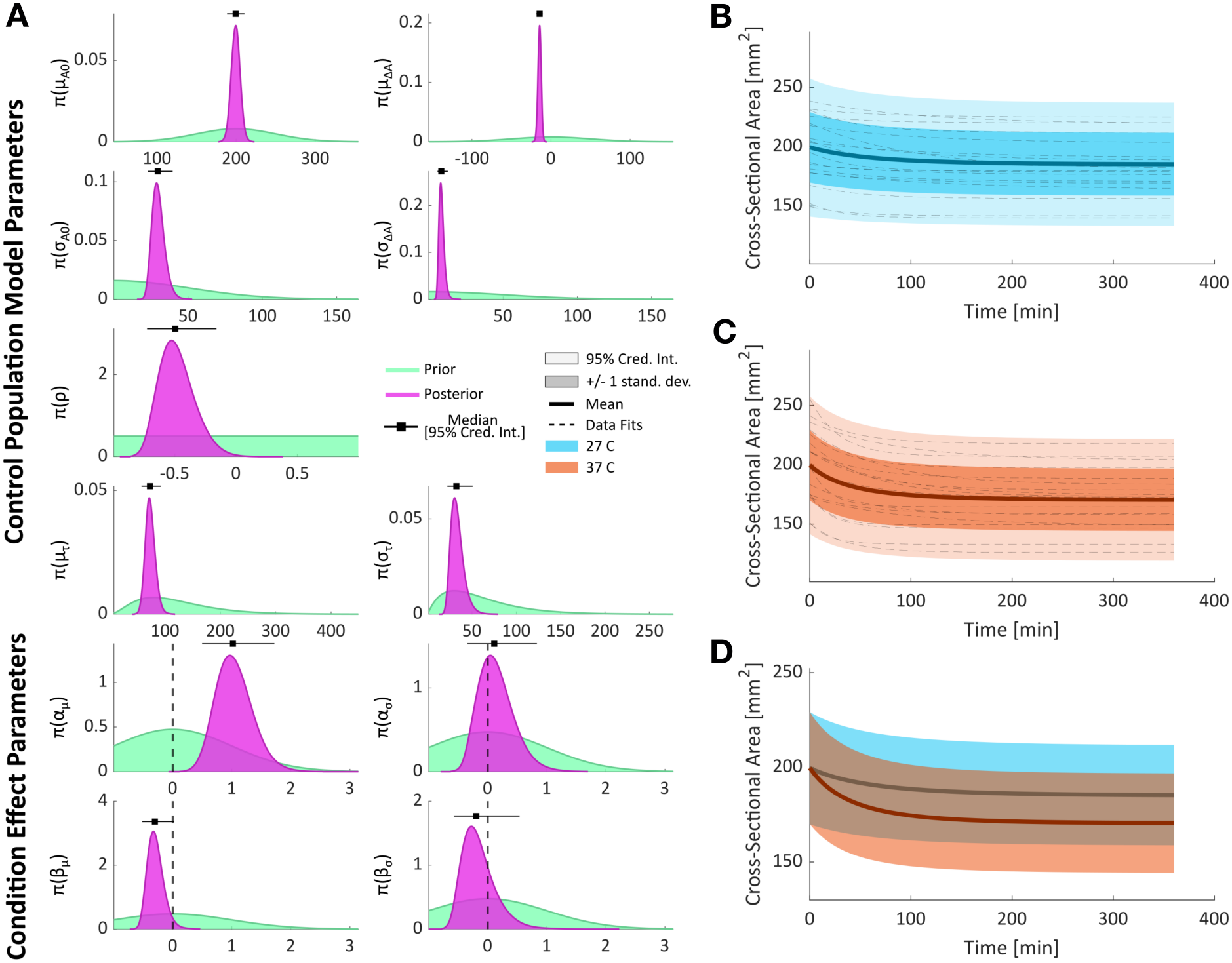
Population modeling of cross-sectional area dynamics. A) Prior and posterior distributions for each fitted parameter. Green curves show the prior distribution and pink curves show the posterior density inferred by SMC, where the y axis represents probability density *π* and x axis width reflects parameter uncertainty. Narrower distributions indicate stronger constraint by the data and shifts relative to the prior reflect parameter updating during inference. Black squares: median, horizontal line: 95% credible interval. For the condition effect parameters, the vertical dashed line is the null effect (no change with temperature). B,C) Median posterior predictive check for the B) 27^*°*^C and C) 37^*°*^ groups cross-sectional area over time of FRESH-printed alginate samples (solid line: mean, dark shaded region: *±*1 standard deviation, light shaded regions: 95% prediction interval, dashed lines: individual dynamical models fit to experimental data). The median posterior model successfully encapsulates all of the observed data in both groups. D) Comparison of 27^*°*^ and 37^*°*^ trajectories (mean and std) as a result of temperature condition effect parameters *α* and *β*. Note: the mean and standard deviation regions in D are the same as shown in B and C.

**Figure 9.**
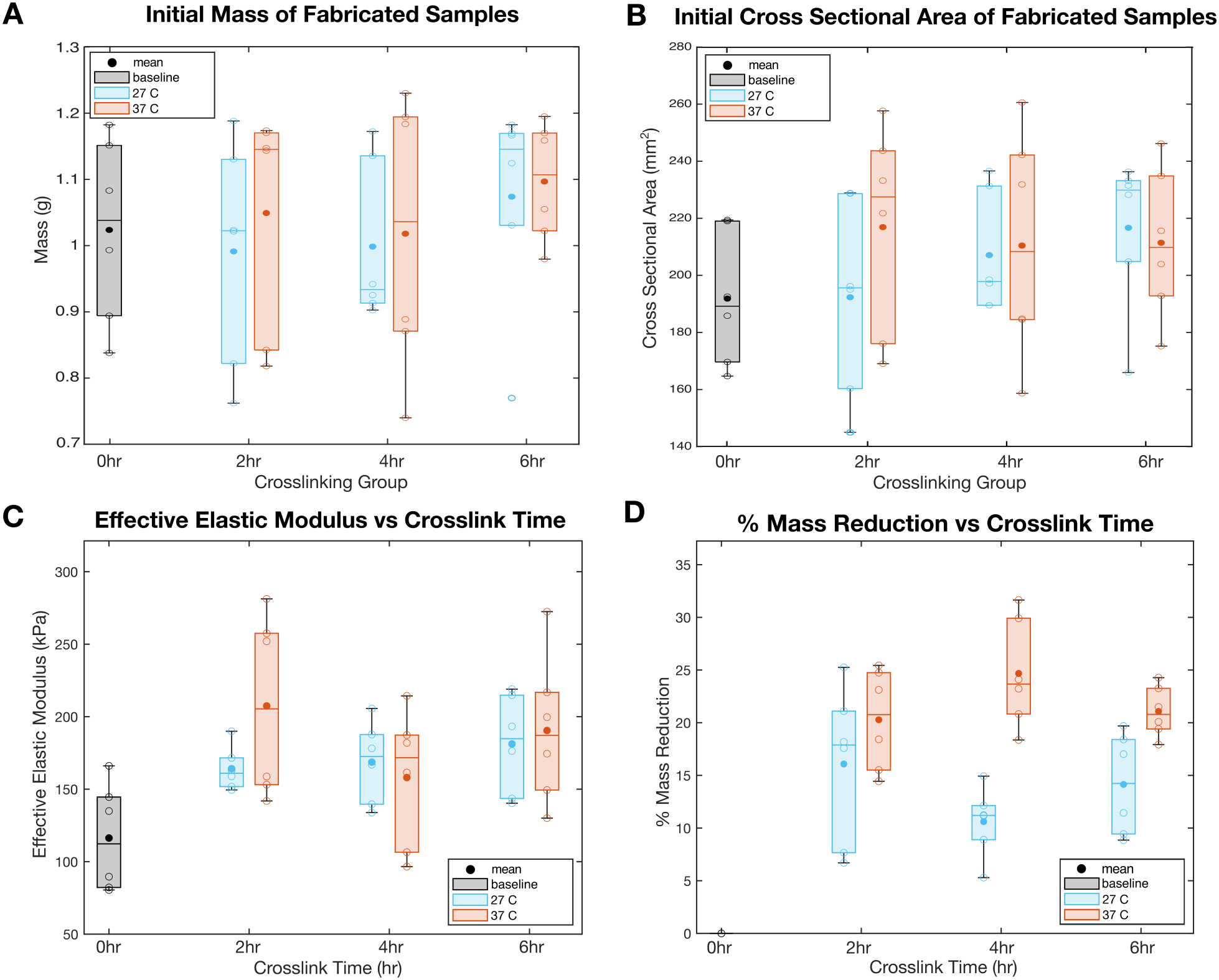
Effects of controlled cross-linking conditions on alginate material properties. A) Initial mass of alginate FRESH-printed constructs with respect to discrete cross-linking time and temperature groups (N=6), immediately after baseline cross-linking (*t* = 0). B) Initial cross-sectional area of FRESH-printed constructs with respect to discrete cross-linking time and temperature groups (N=6). C) Effective elastic modulus of alginate FRESH-printed constructs with respect to discrete cross-linking time and temperature groups (N=6). D) Percent mass reduction experienced by FRESH-printed alginate constructs with respect to discrete cross-linking time and temperature groups (N=6).

Compared to the initial starting condition geometry, there was relatively lower variability observed in the contraction magnitude (*σ*_Δ*A*_ = 8.45 [6.18, 12.36] mm^2^). This variability being smaller than that associated with the initial geometry suggests that, although individual samples begin with substantially different cross-sectional areas, the contraction process itself is comparatively consistent across the population. On average, this contraction (*µ*_Δ*A*_ = *−*14.48 [−18.20, −11.05] mm^2^) represents a measurable portion of the initial CSA (~7% average change), confirming that the ionic cross-linking may meaningfully change the geometry of parts after fabrication.

Additionally, the correlation coefficient *ρ* was strongly negative, *ρ* = *−*0.50 [−0.71, −0.20], and excluded zero across its 95% credible interval. This provides evidence for a relationship between the initial construct geometry and the subsequent contraction behavior. Because Δ*A* is defined as a negative change in area, this result indicates that constructs with larger initial cross-sectional areas tended to experience larger absolute reductions in area during ionic cross-linking. This strong correlation suggests that cross-linking may be causing cross-sectional area changes by a random stretch factor, rather than a random additive change. If the final CSA *A*_*final,i*_ *≈ A*_0,*i*_*λ*_*i*_, for some stretch factor *λ*_*i*_ *≈ µ*_*λ*_ +*ϵ*(*σ*_*λ*_) (*λ*_*i*_ *<* 1), then Δ*A*_*i*_ = *A*_*final,i*_ *− A*_0,*i*_ = (*λ*_*i*_ *−* 1)*A*_0,*i*_ and we would expect Δ*A*_*i*_ and *A*_0,*i*_ to be strongly negatively correlated, as *A*_0,*i*_ would be a direct factor in determining Δ*A*_*i*_. By re-parameterizing the dynamical model in terms of these stretch factors, one could investigate correlations between the absolute size of the polymer network and the fractional size change it undergoes during cross-linking, which may provide interesting insights into the chemo-mechanics of the process.

The kinetic parameters further characterize the contraction process. The mean contraction timescale was estimated as *µ*_*τ*_ = 73.05 min [59.25, 90.36], indicating that the majority of geometric contraction occurred within approximately the first hour of ionic cross-linking. The variability in these kinetics, however, was substantial (*σ*_*τ*_ = 32.30 [23.45, 48.69] min), demonstrating that samples approached their steady-state geometries at different rates despite exhibiting similar first-order behavior. If we specify that steady state has been achieved when a sample has gone through 95% of its change (*A*(*t*_*steady*_) *− A*_0_ = 0.95Δ*A*), this level of kinetics variability suggests that the time to reach steady state will vary greatly (*t*_*steady*_ = 3.63 [1.19, 7.40] hours; median [95% prediction interval]). If we require a tighter definition of steady state, this variability will become even larger as the kinetics slow down. It also suggests that while a typical sample may reach steady state within 4 hours, if a particular application were to require all samples to have achieved steady state, one would have to wait much longer to ensure this.

The temperature-effect parameters provide further insight into how cross-linking conditions influence the contraction process. The positive value of *α*_*µ*_ (1.02 [0.56, 1.66]) indicates that elevated temperature roughly doubles the magnitude of contraction, leading to much greater deviations in the final geometry. Additionally, the negative value of *β*_*µ*_ (−0.30 [−0.49, −0.02]) indicates a much faster approach to steady state (*t*_*steady*_(37^*°*^*C*) = 2.53 [0.66, 5.60]). In contrast, the credible intervals for both *α*_*σ*_ and *β*_*σ*_ overlapped zero, suggesting little evidence that temperature meaningfully altered the variability in either contraction magnitude or contraction kinetics.

The resulting median posterior model provides probabilistic predictions of construct geometry throughout cross-linking. For a FRESH-printed alginate construct with a baseline cross-sectional area of approximately 200 mm^2^, the model predicts a mean reduction in area of approximately 14 mm^2^ under the baseline temperature condition, corresponding to a final steady-state cross-sectional area near 185 mm^2^ (Figure 8B). Elevated-temperature cross-linking approximately doubles the expected contraction magnitude through the positive temperature effect parameter *α*_*µ*_, resulting in a predicted steady-state area near 170 mm^2^ (Figure 8C). Importantly, these predictions are accompanied by confidence bounds arising from the inferred population variability, enabling estimation of not only the expected final geometry, but also the range of geometric outcomes that may be observed both within groups of identically fabricated constructs as well as across groups subject to varying fabrication conditions (Figure 8D).

### Effect of cross-linking conditions on stiffness and mass reduction

#### Group-level analysis of discrete data fails to capture dynamic or population-level behavior

The material variability identified through ^1^H-NMR analysis was also evident in the physical characteristics of the printed constructs prior to extended cross-linking. Although samples within experimental groups were fabricated using identical printing and post-processing protocols, appreciable variability was observed in both initial mass (Figure 9A) and initial cross-sectional area (Figure 9B) immediately following slurry liquefaction. The broad distributions indicate that specimen-to-specimen differences were already present at the start of the experiment. Because the effective elastic modulus was calculated with respect to the cross-sectional area of each individual sample, this metric is more agnostic to geometric deviations resulting from manufacturing variability due to the printing process, and therefore provides more direct evidence than initial CSA analysis alone that heterogeneity in samples exists due to material variability in the hydrogel population before subsequent material evolution occurs and is expected to contribute to variability in the final material response.

This population-level variability was also reflected in the measured mechanical and swelling behavior of the hydrogels. Baseline-cross-linked samples exhibited a mean effective elastic modulus of 120 *±* 40 kPa, reflecting the mechanical behavior of the baseline cross-linked polymer network (Figure 9C). After extended cross-linking for 6 hours, samples cross-linked at 27^*°*^C reached a mean effective elastic modulus of 180 *±* 30 kPa, while those cross-linked at 37^*°*^C reached 190 *±* 50 kPa. In the swelling experiment, samples cross-linked for 6 hours at 27^*°*^C experienced a mean mass reduction of 14 *±* 5%, while those cross-linked at 37^*°*^C experienced a mean mass reduction of 21 *±* 2% (Figure 9D). Detailed results for each experimental time and temperature group can be found in Supporting Information Table SS4.

Although differences in mean values were observed between experimental time and temperature groups, the large standard deviations relative to these differences make it difficult to confidently distinguish material behavior based on group-level summary statistics alone. More importantly, reporting only the mean and standard deviation provides little information regarding the underlying dynamic processes responsible for the observed material response. To investigate whether the observed responses were consistent with the proposed first-order constitutive model (Eqn. 4), the group-level experimental data for both effective elastic modulus *E*_*eff*_ and mass reduction *M* were fit directly to

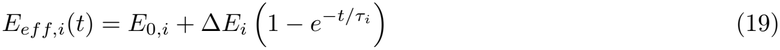

and

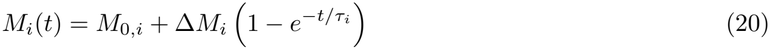

respectively, but resulted in low coefficients of determination 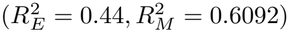 even within 27^*°*^*C* temperature conditions (Figure SS3). However, poor group-level fits cannot be used to determine whether the first-order model is inappropriate. Unlike the cross-sectional area measurements, which consisted of repeated observations collected from the same specimen over time, both effective elastic modulus and mass reduction were obtained through destructive or sparse endpoint assays. Consequently, individual samples could not be tracked throughout the cross-linking process, and the absence of longitudinal measurements prevents direct verification of the behavior of individual samples and obscures the relationship between dynamic material behavior and population-level variability.

Therefore, the population-based modeling framework is required to infer both the likely dynamic trajectories and the uncertainty associated with biological variability. Furthermore, because individual sample trajectories are unobservable, dynamical model parameters for each sample cannot be estimated, and the likelihood of the dynamic parameters cannot be directly calculated. To overcome this limitation, the calibration routine of the population modeling framework was supplemented with Synthetic Likelihood (SL) methods capable of inferring first-order population dynamics from sparse and destructive datasets.

#### Population modeling of effective elastic modulus enables prediction of population-level stiffness evolution and variability

To overcome the uncertainty in steady-state effective elastic modulus revealed by the experimental data, we calibrated our generative stiffness model using a Synthetic Likelihood–Sequential Monte Carlo (SL–SMC) approach. The resulting posterior distributions (Figure 10A) yielded parameter estimates (Table 3) that were substituted into the model to generate median posterior-predictive stiffness trajectories for both temperature conditions (Figure 10B and C). These posterior simulations enabled estimation of steady-state stiffness behavior and the expected spread of modulus values arising from intrinsic material variability and comparative cross-linking conditions (Figure 10D). Joint posteriors, probability density contours, and convergence over SMC generations are provided in the Supporting Information Figure S5.

**Table 3:** Posterior parameter outputs from converged Effective Elastic Modulus population model.

| parameter | median | 95% CI |
|---|---|---|
| μ<sub>**E<sub>0</sub>**</sub> (kPa) | 124.162 | [96.227,156.021] |
| σ<sub>**E<sub>0</sub>**</sub> (kPa) | 33.343 | [21.030,59.535] |
| μ<sub>**ΔE**</sub> (kPa) | 52.359 | [16.292,87.697] |
| σ<sub>**ΔE**</sub> (kPa) | 35.708 | [4.753,77.506] |
| α<sub>μ</sub> (unitless) | 0.255 | [ -0.394,1.252 ] |
| α<sub>σ</sub> (unitless) | 0.724 | [ -0.900,2.221 ] |
| ρ | -0.662 | [-0.989,0.760] |
| μ<sub>τ</sub> (min) | 89.720 | [18.883,240.486] |
| σ<sub>τ</sub> (min) | 51.262 | [7.632,176.749] |
| β<sub>μ</sub> (unitless) | -0.342 | [ -0.962,1.496 ] |
| β<sub>σ</sub> (unitless) | 0.442 | [ -0.880,2.313 ] |

**Figure 10.**
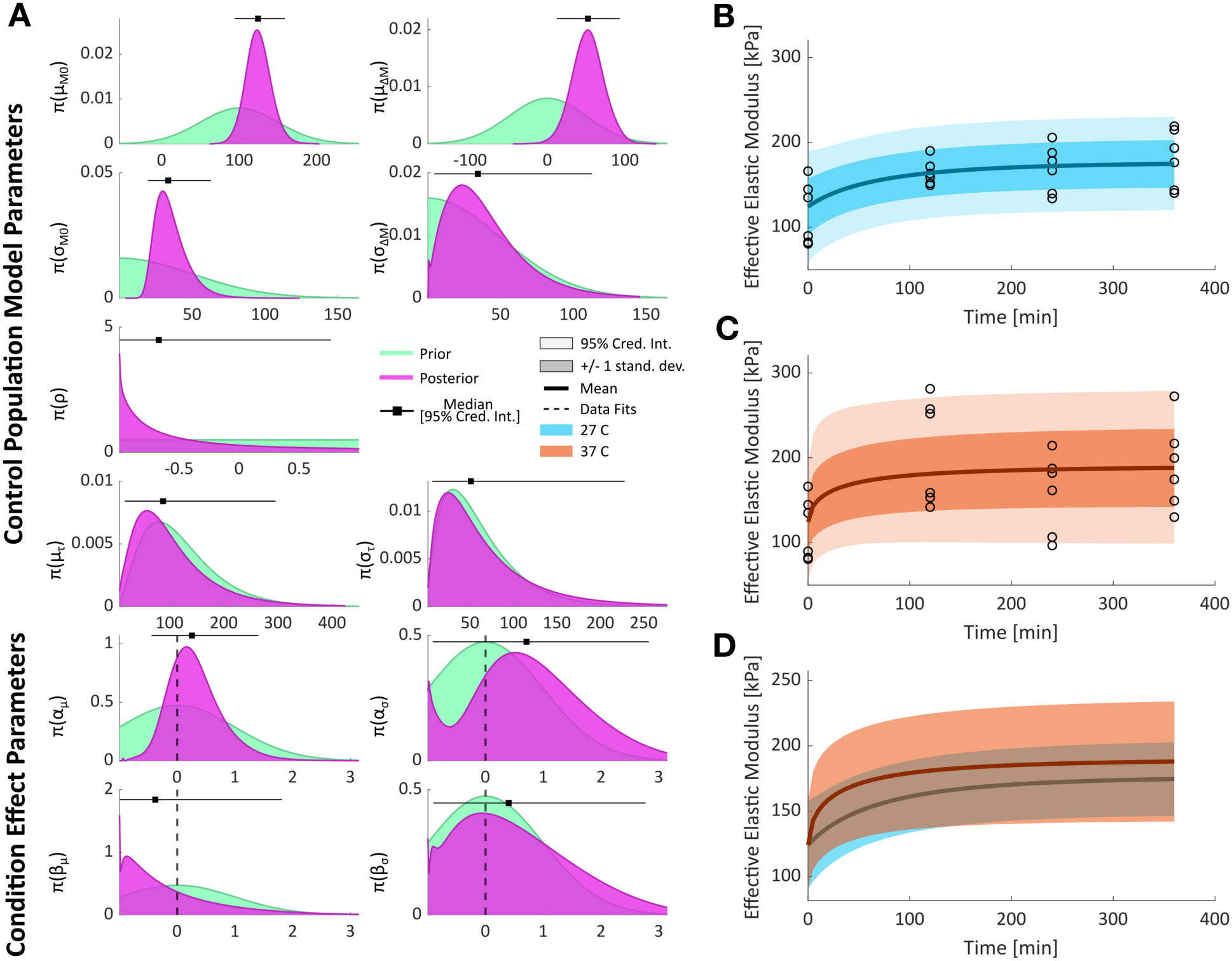
Population modeling of effective elastic modulus dynamics. A) Prior and posterior distributions for each fitted parameter. Green curves show the prior distribution and pink curves show the posterior density inferred by SMC, where the y axis represents probability density *π* and x axis width reflects parameter uncertainty. Narrower distributions indicate stronger constraint by the data and shifts relative to the prior reflect parameter updating during inference. Black squares: median, horizontal line: 95% credible interval. For the condition effect parameters, the vertical dashed line is the null effect (no change with temperature). B,C) Median posterior predictive check for the B) 27^*°*^C and C) 37^*°*^ groups of the effective elastic modulus over time of FRESH-printed alginate samples (solid line: mean, dark shaded region: *±*1 standard deviation, light shaded regions: 95% prediction interval, circles: individual end-point data of the effective elastic modulus). The median posterior model successfully encapsulates all the observed data in the baseline temperature group. Some data in the 37^*°*^C group fall outside of the 95% prediction interval, suggesting, the distribution may have heavier tails than assumed with our population model. D) Comparison of 27^*°*^ and 37^*°*^ trajectories (mean and std) as a result of temperature condition effect parameters *α* and *β*. Note: the mean and standard deviation regions in D are the same as shown in B and C.

The posterior distributions revealed several important features of the underlying material variability. First, the relatively large variability in the baseline-cross-linked stiffness parameter, reflected in the high value of 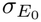 (33.34 [21.03, 59.54] kPa) relative to the mean 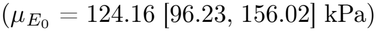 suggests that even before changes induced by cross-inking, monomer composition propagates to stiffness variability. The posterior median for the stiffening parameter was positive (*µ*_Δ*E*_ = 52.36 [16.29, 87.70] kPa), reflecting that extended ionic cross-linking significantly increased the effective elastic modulus. The posterior median for the variability in stiffening (*σ*_Δ*E*_ = 35.71 [4.75, 77.51] kPa) suggests that the degree of mechanical reinforcement achieved during CaCl_2_ exposure may change substantially across samples. However, the lack of dynamical data, and the subsequent correlation between the stiffening and time constant terms, left our estimate of this effect uncertain, reflected in its broad posterior credible interval.

The posterior correlation parameter *ρ* (−0.66 [−0.99, 0.76]) was broadly distributed, indicating that no reliable relationship could be established between the stiffness of the baseline-cross-linked material and the stiffness achieved after full ionic cross-linking. Although the posterior median suggests a potential tendency for initially softer samples to experience larger stiffness gains, the broad credible interval indicates that the available data are insufficient to confidently distinguish this behavior from a positive correlation or the absence of correlation altogether.

The kinetic parameter *τ* provides additional insight into the progression toward steady-state stiffness. The posterior median for *τ* was approximately 90 min, but displayed a broad credible interval (89.72 [18.88, 240.49] min). The posterior for *σ*_*τ*_ was similarly broad (51.26 [7.63, 176.75] min). However, the high uncertainty in both *µ*_*τ*_ and *σ*_*τ*_ suggests that the available stiffness measurements primarily enable prediction of the final material state rather than the exact pathway used to reach it. This uncertainty is expected given that effective elastic modulus measurements were obtained only at discrete two-hour intervals and individual samples could not be tracked longitudinally. As a result, multiple combinations of stiffening magnitude and timescale can produce similarly plausible fits to the observed endpoint measurements. Application of the framework to continuously sampled geometric datasets, particularly with non-destructive mechanical characterization methods, would provide much stronger constraints on the kinetic parameters.

The temperature-effect parameters provide only limited evidence that cross-linking temperature influences the stiffness response. Although the posterior medians suggest increased stiffening (*α*_*µ*_ = 0.26) and faster kinetics (*β*_*µ*_ = *−*0.34) at elevated temperature, the credible intervals for all four temperature-effect parameters (*α*_*µ*_, *α*_*σ*_, *β*_*µ*_, and *β*_*σ*_) encompassed zero. As a result, the available data do not support a definitive conclusion that temperature substantially alters either the magnitude or variability of stiffness development. This result is consistent with the broad experimental overlap observed between the temperature groups (Figure 10D). More confident conclusions may be drawn with additional longitudinally collected data.

#### Population modeling of mass reduction enables prediction of population-level polymer contraction and variability

We calibrated our model describing mass reduction using the same Synthetic Likelihood–Sequential Monte Carlo (SL–SMC) approach as with effective elastic modulus. The resulting posterior distributions in Figure 11A yielded parameter estimates (Table 4) that were substituted into the model to generate temperature-dependent posterior-predictive mass reduction trajectories (Figure 11B and C). These posterior simulations allowed us to estimate and compare the mass reduction experienced by printed samples undergoing polymer contraction in different cross-linking conditions, as well as visualize the population-level variability in the polymer network behavior (Figure 11D). Joint posteriors, probability density contours, and convergence over SMC generations are provided in the Supporting Information Figure S6.

**Table 4:** Posterior parameter outputs from converged Mass Reduction population model.

| parameter | median | 95% CI |
|---|---|---|
| μ<sub>**M<sub>0</sub>**</sub> (g) | 1.034 | [0.992,1.077] |
| σ<sub>**M<sub>0</sub>**</sub> (g) | 0.124 | [0.105,0.149] |
| μ<sub>**ΔM**</sub> (g) | -0.144 | [-0.185,-0.112] |
| σ<sub>**ΔM**</sub> (g) | 0.011 | [0.001,0.047] |
| α<sub>μ</sub> (unitless) | 0.622 | [ 0.222,1.159 ] |
| α<sub>σ</sub> (unitless) | -0.163 | [ -0.940,1.811 ] |
| ρ | 0.134 | [-0.896,0.9610] |
| μ<sub>τ</sub> (min) | 52.813 | [12.458,146.686] |
| σ<sub>τ</sub> (min) | 46.134 | [5.045,180.887] |
| β<sub>μ</sub> (unitless) | -0.282 | [ -0.985,1.522 ] |
| β<sub>σ</sub> (unitless) | -0.024 | [ -0.944,2.064 ] |

**Figure 11.**
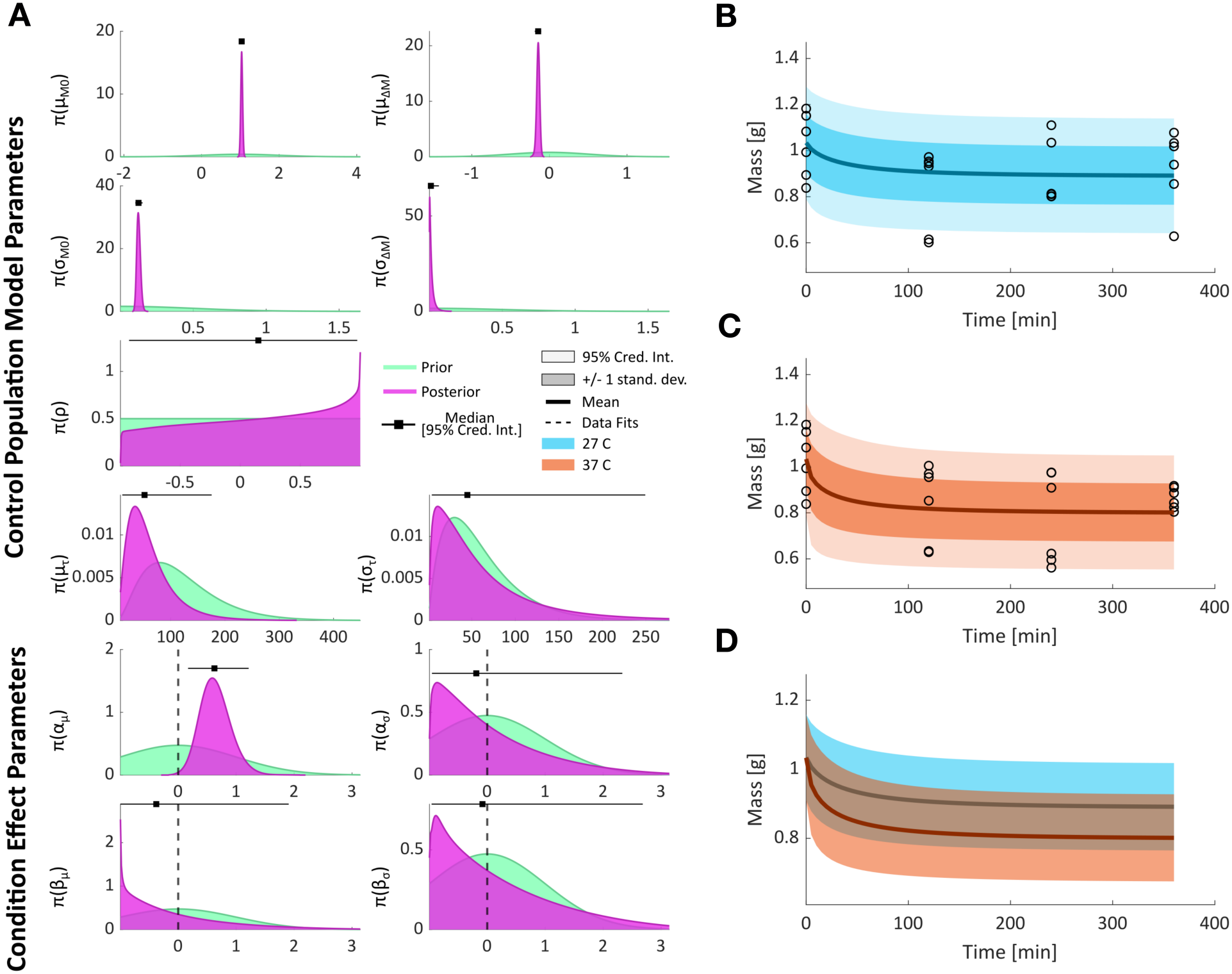
Population modeling of mass reduction dynamics. A) Prior and posterior distributions for each fitted parameter. Green curves show the prior distribution and pink curves show the posterior density inferred by SMC, where the y axis represents probability density *π* and x axis width reflects parameter uncertainty. Narrower distributions indicate stronger constraint by the data and shifts relative to the prior reflect parameter updating during inference. Black squares: median, horizontal line: 95% credible interval. For the condition effect parameters, the vertical dashed line is the null effect (no change with temperature). B,C) Median posterior predictive check for the B) 27^*°*^C and C) 37^*°*^ groups for the mass reduction over time of FRESH-printed alginate samples (solid line: mean, dark shaded region: *±*1 standard deviation, light shaded regions: 95% prediction interval, dashed lines: individual end-point experimental data of the mass). The median posterior model successfully encapsulates all of the observed data in both groups. D) Comparison of 27^*°*^ and 37^*°*^ trajectories (mean and std) as a result of temperature condition effect parameters *α* and *β*. Note: the mean and standard deviation regions in D are the same as shown in B and C.

The posterior distributions indicate that mass reduction during ionic cross-linking was both substantial and comparatively consistent across the hydrogel population. The mean initial construct mass was estimated as 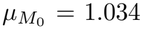 [0.992, 1.077] g, with a corresponding population variability of 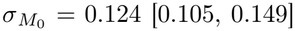 g. This modeled variability is consistent with the spread in initial sample masses observed experimentally (Figure 9A). The posterior median mass-change parameter was strongly negative (*µ*_Δ*M*_ = *−* 0.144 [−0.185, −0.112] g) and the associated variability was comparatively small (*σ*_Δ*M*_ = 0.011 [0.001, 0.047] g), suggesting that although constructs begin with different initial masses, the amount of water expelled due to contraction of the polymer network during cross-linking is relatively consistent between samples.

The posterior correlation parameter remained largely unresolved, with *ρ* = 0.134 [−0.896, 0.961]. The broad credible interval indicates little evidence for a systematic relationship between a construct’s initial mass and the magnitude of mass reduction experienced during cross-linking. The inferred kinetic parameters suggest that mass equilibration occurs relatively rapidly, with a posterior median time constant of *µ*_*τ*_ = 52.81 [12.46, 146.69] min. However, both *µ*_*τ*_ and *σ*_*τ*_ (46.13 [5.05, 180.89] min) exhibited broad credible intervals. As with the stiffness data, the uncertainty in correlation and kinetic parameters reflects the limited temporal information and the lack of longitudinal relationships present in the experimental data.

Among the temperature-effect parameters, only mean in mass change parameter (*α*_*µ*_) showed strong evidence of a non-zero temperature dependence (0.622 [0.222, 1.159]). This result indicates that elevated cross-linking temperature increased the magnitude of mass reduction relative to the baseline condition. In contrast, the credible intervals for *α*_*σ*_, *β*_*µ*_, and *β*_*σ*_ all encompassed zero, suggesting limited evidence that temperature substantially altered either the variability of mass loss or the kinetics of the equilibration process.

### Comparison of material models reveals common mechanisms governing hydrogel variability

Although cross-sectional area, effective elastic modulus, and mass reduction quantify different material responses, considering these metrics together provides a more complete picture of the underlying polymer network evolution during ionic cross-linking. Cross-sectional area directly quantifies geometric fidelity and therefore the dimensional accuracy that can be expected in a manufactured hydrogel component. Effective elastic modulus provides insight into the mechanical integrity of the network and its ability to resist deformation, while mass reduction serves as an indirect measure of polymer network contraction through water expulsion during equilibration. Collectively, these metrics span both macroscopic performance and microscopic network behavior, allowing the effects of cross-linking to be evaluated from complementary perspectives.

A consistent trend across all three models was the presence of substantial variability in the initial state of the hydrogel population. Large posterior median estimates of *σ*_*X*0_ (*X ∈* [*A, E, M*]), reflecting broad distributions for *A*_0_, *E*_0_, and *M*_0_, indicate that appreciable heterogeneity exists immediately following printing and baseline cross-linking. Interestingly, the cross-sectional area and mass-reduction models both indicated that variability in the change term was substantially smaller than variability in the corresponding initialstate parameter. In other words, samples began from different baseline conditions but tended to undergo relatively similar contraction processes once exposed to extended ionic cross-linking. Although the effective elastic modulus model exhibited greater uncertainty in the corresponding stiffening parameter, its posterior distributions remained consistent with this broader pattern. From a manufacturing perspective, this finding suggests that variability in the stock materials and ink composition is a substantial contributor to variability in final printed constructs, and is only compounded and exacerbated further by any variability introduced during the primary FRESH printing or secondary CaCl_2_ cross-linking processes themselves. For example, differences in polymer chain length and shape may influence ink viscosity, which may propagate to differences in how ink flows and spreads when extruded into the slurry, resulting in a wide distribution of initial printed geometries even before slurry release.

The posterior correlation parameters provide additional support for this interpretation. The cross-sectional area model identified a potential negative relationship between initial area and subsequent contraction magnitude, while the effective modulus model exhibited a similarly negative posterior median despite substantially greater uncertainty. Although these observations do not directly demonstrate a common mechanism, they are consistent with the possibility that samples enter the extended cross-linking phase at different stages of network formation. Under such a scenario, constructs that are initially larger or mechanically weaker would not necessarily remain so indefinitely, but could instead experience larger subsequent changes as they converge toward a common equilibrium state shared by the population. This interpretation is further supported by the broad distributions observed for the kinetic parameters across all three material models, suggesting that a meaningful component of the observed heterogeneity may arise from differences in the rate at which samples progress through similar cross-linking states rather than from fundamentally different equilibrium behaviors or cross-linking capacity. The modeling framework could be applied in the future to additional experiments that investigate this concept by comparing the variability in mechanical performance of samples with equivalent contraction magnitude, rather than equivalent cross-link time points. One possible mechanistic explanation for these trends is the coexistence of two distinct classes of network interactions within alginate hydrogels. Prior to ionic cross-linking, material behavior is strongly influenced by physical entanglement and hydrogen-bond-mediated interactions that emerge from chain length, chain architecture, and polymer overlap [19, 23]. These relationships are highly variably across biological species [26] and may be expected to produce differences in the initial geometry, water content, and mechanical properties of printed constructs. In contrast, the stiffening that occurs during CaCl_2_ exposure is governed by ionic cross-linking, a process that is more controllable and uniform [19], and therefore may behave more consistently across samples. The H-NMR composition data reflects this. The three alginate batches exhibited only modest differences in their overall M/G ratios (0.5530, 0.6034, 0.9048), which correspond to the density potential of ionically cross-linkable sites. Conversely, they showed much larger variability in their MM/GG doublet ratios (1.6838, 0.0437, 0.8144), which reflect the presence of repeating blocks that indicate high molecular weights, long polymer chains, and potentially impact the degree of physical entanglement of the network [19, 26]. While the present study cannot directly separate the contributions of these two network structures, the collective posterior trends are consistent with a system in which physical entanglement drives much of the initial heterogeneity while ionic cross-linking acts as a more uniform network-strengthening mechanism.

The temperature-effect parameters provide the clearest process-level insight. Across both the cross-sectional area and mass-reduction models, elevated temperature consistently and significantly increased the magnitude of contraction, indicating greater geometric deviation and greater water loss during cross-linking. However, these changes were not accompanied by a similarly supported increase in the effective elastic modulus. The posterior median model associated with stiffness showed substantial overlap between temperature conditions, providing little evidence that elevated temperature produced a mechanically advantageous final material state. If anything, the broader stiffness distributions observed at elevated temperature may suggest increased heterogeneity in network development, although this possibility would require more direct investigation in future work. Collectively, these results indicate that higher cross-linking temperatures primarily accelerate and amplify contraction while providing no clear mechanical benefit. For applications in which geometric fidelity, dimensional accuracy, and repeatability are important, the results therefore support the use of lower-temperature extended cross-linking conditions. Based on shared posterior parameter predictions for the kinetic time constant *τ*, this extended cross-linking duration should exceed 8 hours to confidently ensure complete cross-linking of all samples while accounting for the variability in their approach to steady-state.

## Conclusion

In this work, we developed a framework combining investigation of polymer network variability, controlled cross-linking experiments, and a probabilistic modeling framework to characterize biologically derived hydrogels. Dynamic datasets, such as cross-sectional area evolution, were incorporated directly through Sequential Monte Carlo estimation, while sparse endpoint measurements were analyzed through extension with Synthetic Likelihood methods. Together, these approaches enable inference of population-level dynamics, providing quantitative insight into material variability, cross-linking behavior, and expected steady-state performance distributions, even when repeated measurements of individual samples are not experimentally feasible.

Using sodium alginate as a case study, we found that substantial variability was present prior to extended ionic cross-linking and contributed significantly to the final distributions of geometric and mechanical properties. By applying the framework across geometric, mechanical, and mass-transfer measurements, we identified several shared trends that were not readily apparent from conventional Frequentist analysis of group means alone, including impacts of organic material variability on the repeatability of final FRESH-printed constructs, possible underlying mechanistic explanations for cross-linking behavior, and suggestions for the reduction of variability during manufacturing. The results suggest that variability introduced during fabrication and baseline cross-linking may be a more important determinant of final material performance than variability introduced during secondary ionic cross-linking.

Importantly, the framework is not specific to alginate or to the properties experimentally measured in this work. The application of the generalizable framework to multiple metrics demonstrates its flexibility with materials and settings where direct measurement is noisy, incomplete, or experimentally burdensome. This method complements and extends prior efforts to model hydrogel mechanics with a streamlined way to approximate bulk, steady-state-relevant behavior that focuses on the emergent material properties most relevant to device and robot design with cross-linkable polymers (initial conditions, magnitude gain, and the rate of approach to equilibrium), without requiring detailed assumptions about polymer network structure. The SL-SMC inference not only provides estimates of the average response of a material property, but allows for quantitative prediction of part variability and indirect estimates of population-level performance variability.

Further work should validate these predictions through longer-duration cross-linking studies with improved temporal resolution in early stages where kinetic uncertainty is greatest, and by incorporating non-destructive mechanical characterization methods that permit repeated measurements of the same sample over time. Applying the framework to additional material properties, such as direct measurement of polymer network behavior, viscoelastic response, or degradation, may further expand its utility. In addition, its application to alternative fabrication strategies could enable the isolation and characterization of manufacturing variability and fidelity. More broadly, extending the framework to other hydrogel chemistries or biologically derived tissues will further help establish generalizable, geometry-agnostic strategies for characterizing soft, organic materials, and ultimately support more reliable, tunable design and manufacturing of soft robotic actuators, tissue-mimetic constructs, and biodegradable devices.

## Acknowledgements

The authors thank funding support from NSF DBI 2015317 - NSF/CIHR/ DFG/FRQ/UKRI-MRC Next Generation Networks for Neuroscience Program, and from the Carnegie Mellon University Department of Mechanical Engineering through a Collaborative Grand Challenge award. ASW was supported by a Fellowship from the National Consortium for Graduate Degrees for Minorities in Engineering and Science (GEM), the Steinbrenner Institute, and funding award no. HQ00342110020 from the National Defense Education Program. MJB was supported by the NSF Graduate Research Fellowship Program under Grant No. DGE2140739 and the Carnegie Mellon University College of Engineering Presidential Fellowship. Any opinions, findings, conclusions, or recommendations expressed in this material are those of the authors and do not necessarily reflect the views of the National Science Foundation. The authors also thank Karl Schulz of the Mellon Institute Noonan Lab for access and training regarding the Department of Chemistry NMR Center. Bioprinter graphic courtesy of Shiwarski Tissue Engineering Lab Open Source Designs. Remaining equipment icons and graphics provided by NIAID NIH BioArt Source (bioart.niaid.nih.gov/bioart/) and Servier Medical Art (https://smart.servier.com), licensed under CC BY 4.0 (https://creativecommons.org/licenses/by/4.0/).

## Use of Generative AI Statement

Generative AI tools (Microsoft Copilot) were used in the debugging of the SMC Toolbox validation scripts, in the research of the synthetic likelihood approaches, and to improve the language, readability, and organization of the authors’ own text. All code generated was reviewed and validated by the authors. Grammarly was used to improve the grammar of the text but was not used to synthesize new text. Any changes and suggestions made by the AI editing tools were subsequently reviewed by the authors. The authors take full responsibility for the publication’s content.

## Data Availability

The data underlying this study are openly available on GitHub at [CMU-BORG/Hydrogel-Characterization-Framework] and archived on Zenodo at [doi: 10.5281/zenodo.24538830]. In addition, the Sequential Monte Carlo toolbox and Population Model are available through GitHub. Instructions for the use of either tool are available in the online documentation.

## Supporting Information

The following files are available free of charge.

- Supporting Information: Additional experimental methods and materials, including the Calculation of alginate monomer and doublet ratios from ^1^H-NMR spectroscopy, ^1^H-NMR spectra for all samples, Dynamic raw data from cross-linking experiments by experimental group, Discrete raw data from cross-linking experiments by experimental group, and Detailed methods for population modeling of hydrogel properties.

## Additional Experimental Methods and Results

### Calculation of alginate monomer and doublet ratios from ^1^H-NMR spectroscopy

Methods for calculating monomer ratios were adapted from previous literature [26, 28, 37]. Briefly, peaks corresponding to Guluronic anomeric proton residues G1, Mannuronic acid-dominated blocks (G[G5]M, M[G5]M, [M1]G, or [M1]M), and Guluronic acid-dominated blocks (G[G5]G) were identified and assigned in the downfield spectra region. TopSpin’s internal deconvolution algorithm [38, 39] was used to disentangle overlapping peaks and approximate the integrals of each area. The integrals represent the frequency of G and M residues such that

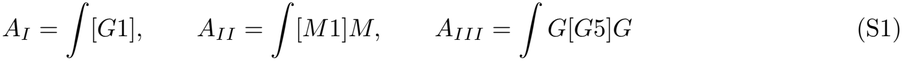

where *A*_*I*_ is the occurrence of G residues, *A*_*II*_ is the occurrence of M blocks, and *A*_*III*_ is the occurrence of G blocks. Given these assignments, the monomer fractions and resulting M/G ratio can be calculated given

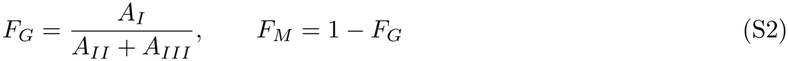

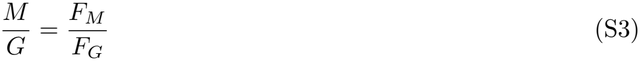

The M/G ratio was used as a quantifiable metric for the relative presence of each type of monomer in the alginate stock material, and as an indication of its likelihood to cross-link fully in a predictable way. A higher ratio indicates a dominance of M monomers, resulting in less uniform cross-linking. Furthermore, it is similarly useful to additionally determine the monomer doublet fractions and ratio as an indication of intact repeating blocks of monomers, which can be calculated given

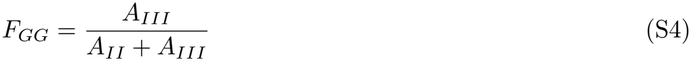

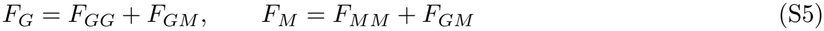

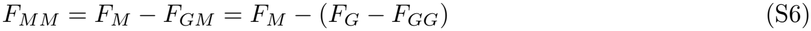

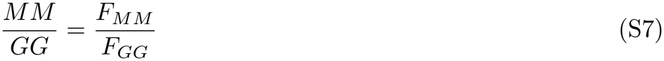

### ^1^H-NMR spectra for all samples

^1^H-NMR spectra for all (N = 3) tested samples can be found in Figure S1. The resulting monomer peak locations and integral values are detailed in Table S1 and Table S2, respectively.

**Figure S1:**
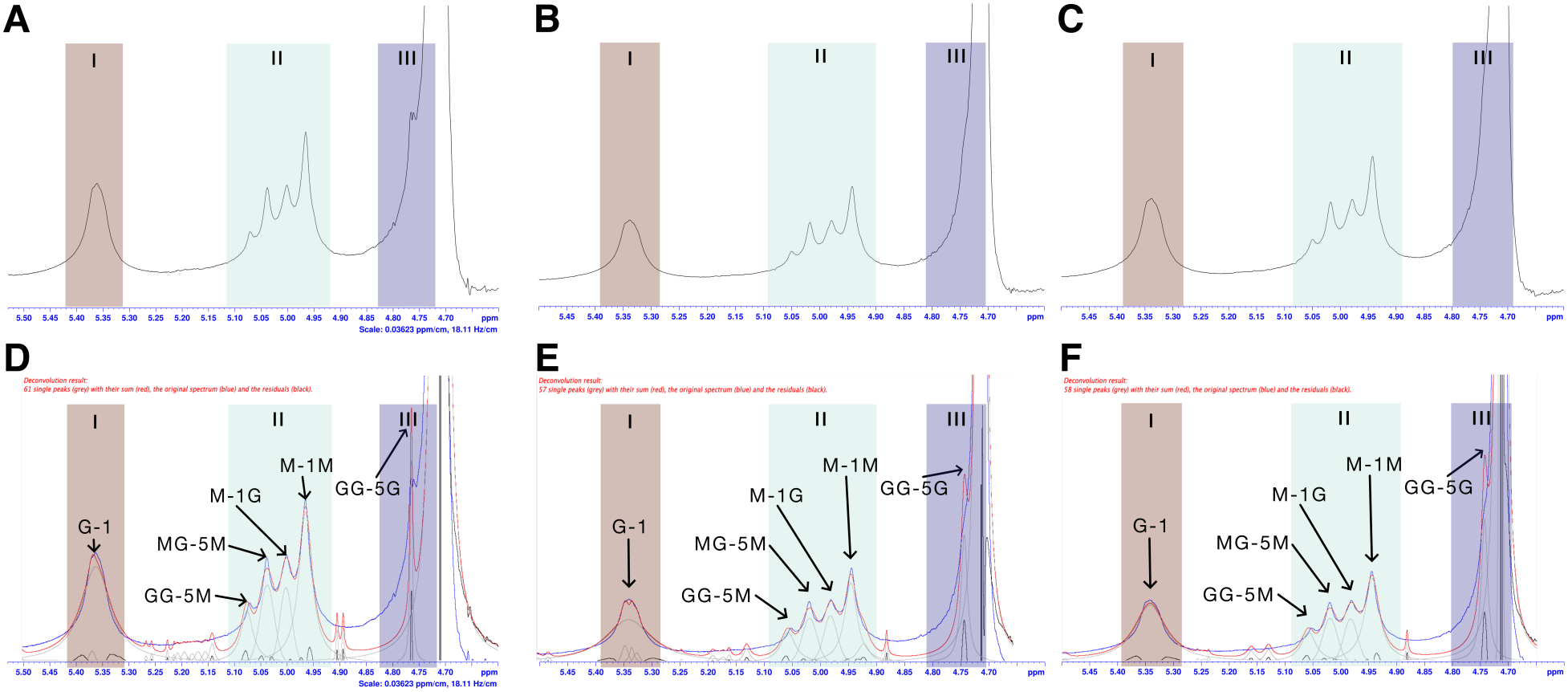
Results of alginate ^1^H-NMR spectral analysis for all samples. A) Sample 1 raw spectra. B) Sample 2 raw spectra. C) Sample 3 raw spectra. D) Sample 1 de-convoluted and assigned peaks. E) Sample 2 de-convoluted and assigned peaks. F) Sample 3 de-convoluted and assigned peaks.

**Table S1:**
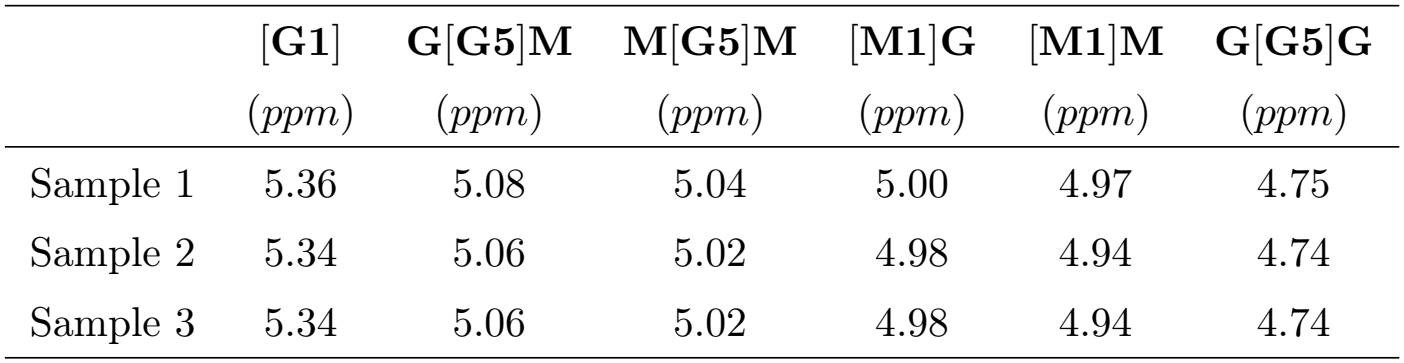
^1^H-NMR spectra peak locations of monomer assignments in analyzed alginate samples.

|  | [**G1**] | **G**[**G5**]**M** | **M**[**G5**]**M** | [**M1**]**G** | [**M1**]**M** | **G**[**G5**]**G** |
|---|---|---|---|---|---|---|
|  | (*ppm*) | (*ppm*) | (*ppm*) | (*ppm*) | (*ppm*) | (*ppm*) |
| Sample 1 | 5.36 | 5.08 | 5.04 | 5.00 | 4.97 | 4.75 |
| Sample 2 | 5.34 | 5.06 | 5.02 | 4.98 | 4.94 | 4.74 |
| Sample 3 | 5.34 | 5.06 | 5.02 | 4.98 | 4.94 | 4.74 |

**Table S2:**
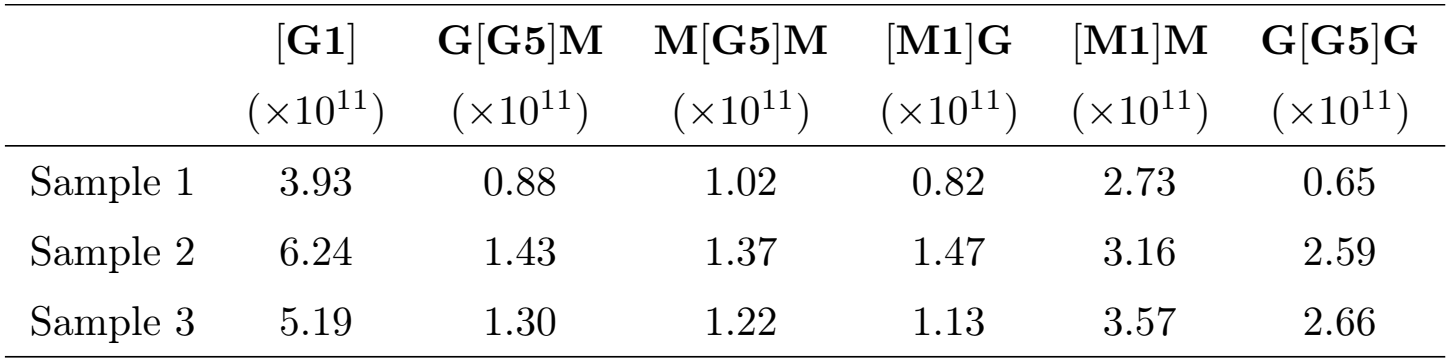
^1^H-NMR spectra integrals of monomer assignments in analyzed alginate samples.

|  | [**G1**]<br>(×10<sup>11</sup>) | **G**[**G5**]**M**<br>(×10<sup>11</sup>) | **M**[**G5**]**M**<br>(×10<sup>11</sup>) | [**M1**]**G**<br>(×10<sup>11</sup>) | [**M1**]**M**<br>(×10<sup>11</sup>) | **G**[**G5**]**G**<br>(×10<sup>11</sup>) |
|---|---|---|---|---|---|---|
| Sample 1 | 3.93 | 0.88 | 1.02 | 0.82 | 2.73 | 0.65 |
| Sample 2 | 6.24 | 1.43 | 1.37 | 1.47 | 3.16 | 2.59 |
| Sample 3 | 5.19 | 1.30 | 1.22 | 1.13 | 3.57 | 2.66 |

### Dynamic raw data from cross-linking experiments by experimental group

To verify that sample-to-sample variability and not ill-fitting model behavior was responsible for the lower *R*^2^ values of each time group, we independently fit each sample to its own exponential model (Figure S2). The resulting average *R*^2^ values all exceeded 0.89.

**Figure S2:**
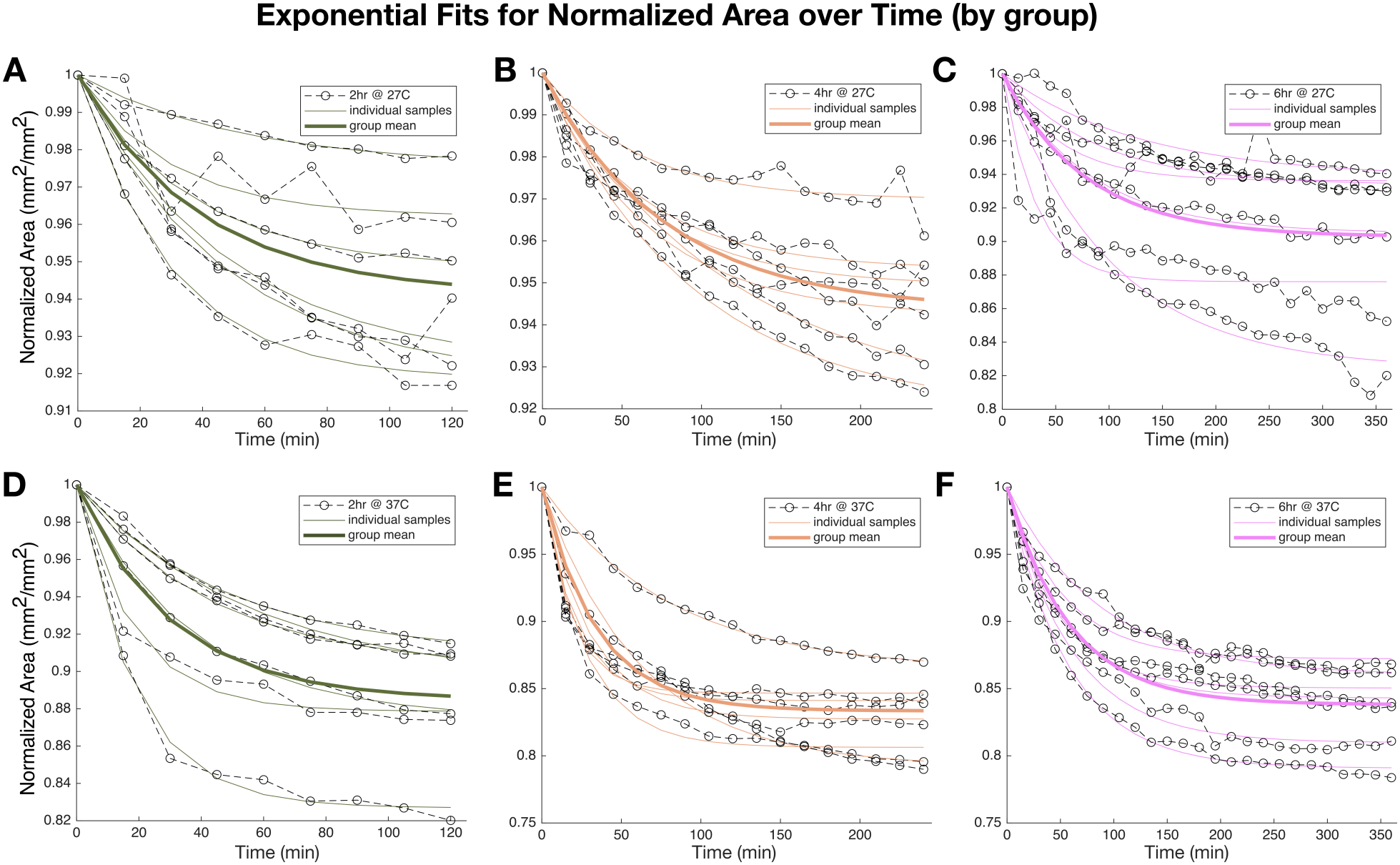
Exponential fits for normalized cross-sectional area over time for each sample in each group. A) Experimental group cross-linked for 2 hr @ 27°C (N=6). B) Experimental group cross-linked for 4 hr @ 27°C (N=6). C) Experimental group cross-linked for 6 hr @ 27°C (N=6). D) Experimental group cross-linked for 2 hr @ 37°C (N=6). E) Experimental group cross-linked for 4 hr @ 37°C (N=6). F) Experimental group cross-linked for 6 hr @ 37°C (N=6). Lines labeled group mean represent models fit to all samples simultaneously.

**Table S3:** Goodness of fit metrics for individual sample and group means in each cross-linking condition group.

|  | cross-link Temp | **2 hr** | **4 hr** | **6 hr** |
|---|---|---|---|---|
| mean *R*<sup>2</sup> of fitted samples | 27°*C* | 0.9378 | 0.9505 | 0.8905 |
|  | 37°*C* | 0.9913 | 0.9767 | 0.9714 |
| *R*<sup>2</sup> of combined group data | 27°*C* | 0.5264 | 0.6806 | 0.3360 |
|  | 37°*C* | 0.5723 | 0.7147 | 0.6943 |

### Discrete raw data from cross-linking experiments by experimental group

The measured mean and standard deviation of stiffness and swelling metrics for each discrete cross-linking time and temperature group are reported in Table S4.

**Table S4:** Mean and std of stiffness and swelling metrics from discrete cross-linking groups.

|  | temp | 0 hr | 2 hr | 4 hr | 6 hr |
|---|---|---|---|---|---|
| Initial Mass (g) | 27°C | 1.02 ± 0.14 | 0.99 ± 0.17 | 1.00 ± 0.12 | 1.07 ±0.16 |
|  | 37°C |  | 1.05 ± 0.17 | 1.02 ± 0.21 | 1.10±0.09 |
| Initial Cross-sectional Area (*mm*<sup>2</sup>) | 27°C | 190 ± 20 | 190 ± 30 | 210 ± 20 | 220 ±30 |
|  | 37°C |  | 220 ± 40 | 210 ± 40 | 210±30 |
| Effective Elastic Modulus (kPa) | 27°C | 120 ± 40 | 170 ± 20 | 170 ± 30 | 180 ±30 |
|  | 37°C |  | 210 ± 60 | 160 ± 50 | 190±50 |
| Mass Reduction (% change) | 27°C | 0 | 16 ± 7 | 10 ±3 | 14 ± 5 |
|  | 37°C |  | 20 ± 5 | 25 ±5 | 21 ±2 |

To evaluate whether the effective elastic modulus and mass reduction followed an exponential approach toward a steady-state value, we fit the experimental data to a first-order exponential model following the generalized form from Eq. 4. The fitted exponential models and R^2^ values for stiffness and swelling data from the 27^*°*^C cross-linking experiments are shown in Figure S3. The poor goodness-of-fit values validate the need for computational modeling to estimate population variability and predict steady-state mechanics as a population rather than the average of individuals.

**Figure S3:**
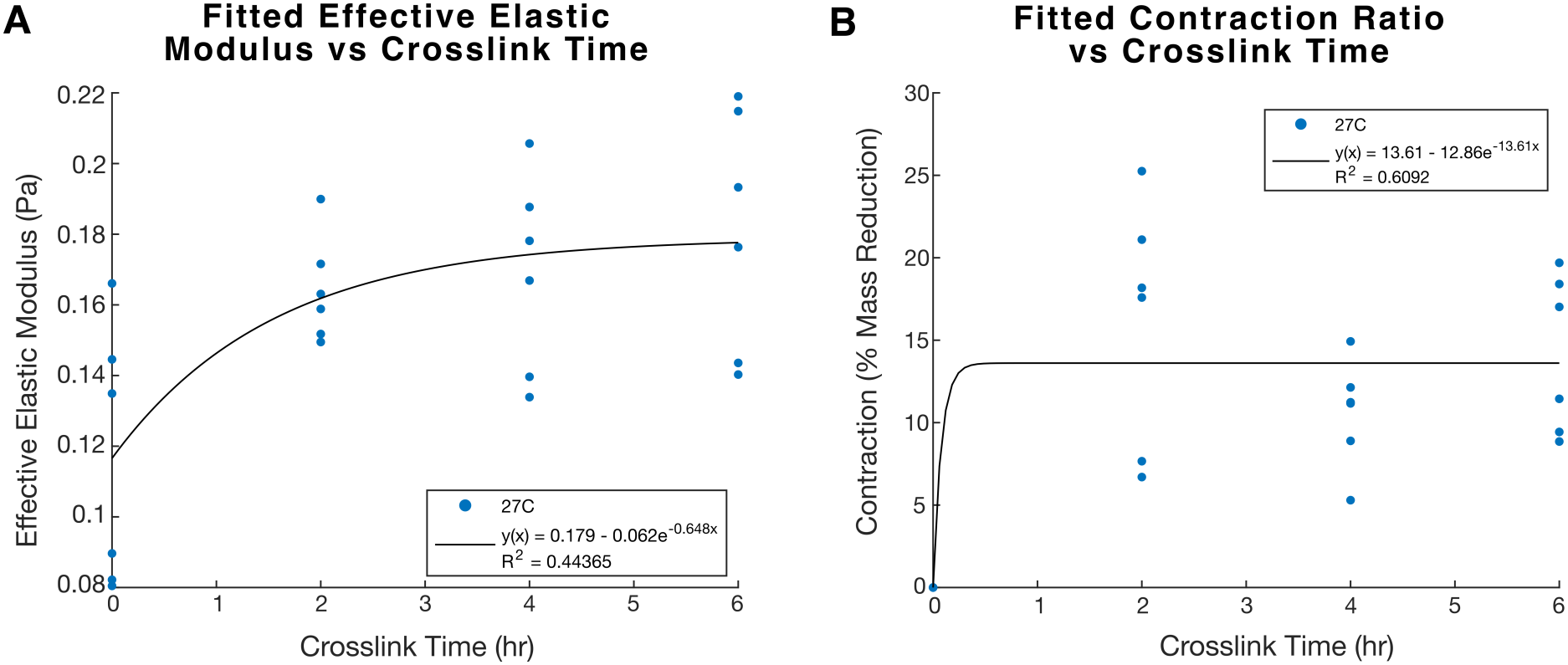
Fitted exponential of data from cross-linking experiments. A) Effective Elastic Modulus experimental data and fitted model. *R*^2^ = 0.44365 B) Percent mass reduction experimental data and fitted model. *R*^2^ = 0.6092

### Summary of Sequential Monte Carlo Methods and Implementation

To estimate the posterior of our population model’s parameters, we utilized a Sequential Monte Carlo method (summarized in Algorithm 1) [41–43]. In this method, we utilized a set of *N* weighted particles 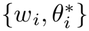, where *w*_*i*_ is the weight of particle 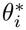) to approximate a distribution of interest. These particles are initially sampled from a set of supplied prior distributions, and the weights are set equal. Then, the weights are updated such that particles that are more probable in the target distribution are weighted more heavily, and those less probable less heavily. This could be used to estimate the weights in the posterior directly. However, if the prior and posterior distributions are very different from each other, this would tend to leave many particles with almost no weight, and a few particles with most of the weight. This problem is referred to as particle degeneracy. To avoid this, SMC methods do not estimate the posterior immediately, but progressively estimate distributions that get closer and closer to the posterior, only estimating the true posterior in the final step. Here, we used geometric bridging, where for a series of *G* values 0 = *γ*_1_ *< γ*_2_ *<* … *< γ*_*G*_ = 1, we estimate the intermediate distributions

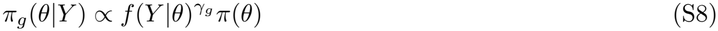

where *f* (*Y*|*θ*) is the likelihood of the dataset *Y*, and *π*(*θ*) is the prior probability [41]. In geometric bridging, *π*_1_(*θ*|*Y*) is exactly the prior, and *π*_*G*_(*θ*|*Y*) is exactly the posterior for which we are searching. In our implementation, we prescribe the values of *γ*_*g*_ as varying linearly from 0 to 1 over a fixed number of generations. However, these can also be adaptively chosen. For synthetic likelihood methods, as utilized for the discrete data sets, the same bridging technique can be used, but with the likelihood replaced by the synthetic likelihood.

At each stage *g*, we update the particle weights for the intermediate distribution *π*_*g*_(*θ*|*Y*) as

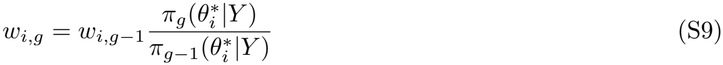

which, for geometric bridging, reduced to

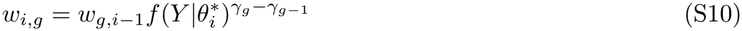

The weights are then normalized so that they all sum to 1. Next, to mitigate the effects of particle degeneracy, we estimate how evenly spread out our weights are by calculating the Effective Sample Size (ESS)

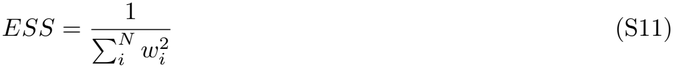

This value ranges from *N*, when all particles have equal weight, to 1, when 1 particle contains the full weight. If the ESS fell below 0.75*N*, the set of particles was resampled using a weighted Systematic Resampling, and the weights were reinitialized to 1*/N*. Finally, to encourage particle mixing and exploration of the parameter space, each particle was independently perturbed using a series of 10 Markov Chain Monte Carlo (MCMC) steps. In each MCMC step, a new proposed particle 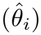 was proposed using a Gaussian random walk as

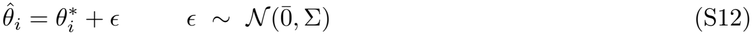

where Σ is a scaled version of the weighted covariance of the particles [43, 48]

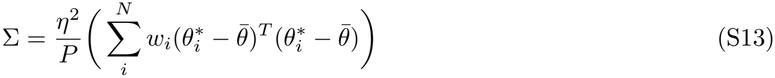

Here, 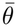 is the weighted mean of the particles, and *P* is the dimensionality of *θ*. This proposed particle is then randomly accepted with probability *p*_*MH*_, where

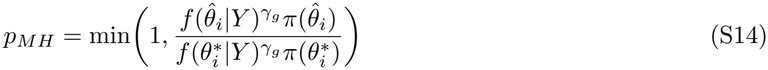

is the Metropolis-Hastings ratio for a symmetric proposal distribution. The value of *η* (the scaling factor applied to the particle covariance) is tuned such that the acceptance rate for a *P >* 1 is between 20% and 40%. The particles that result from the MCMC steps then pass to the next generation. After the end of the final generation, one final systematic resampling is performed to provide an equal weight sample from the posterior. From this final posterior, we can calculate summary metrics, including the posterior median and the 95% credible interval for each parameter, and estimate the posterior probability of a given parameter set using kernel density estimation over the posterior particles.

This algorithm is implemented in the <monospace>SMC_Sampler.m</monospace> class of our <monospace>Sequential Monte Carlo</monospace> MATLAB library. The class is set up to perform SMC estimation of the posterior for an arbitrary statistical model implemented as a MATLAB class (see the GitHub README for requirements of the class). To accelerate the calculation of the posterior particles, both the weight calculations and the MCMC perturbations are calculated in parallel using the Parallel Computing Toolbox. This ability to parallelize is a major advantage of SMC methods over other Bayesian algorithms (e.g., MCMC, Hamiltonian Monte Carlo) which cannot be parallelized in the same way [41, 42]. Finally, if particles contain bounded variables (i.e., elements of the particles are strictly positive or confined between two values), the perturbation step can propose values outside of the allowed range, leading to numerical issues. To avoid this, we implement the ability to use parameter transformations on particular elements of the particle. Specifically, we implement log transformations for strictly positive values and logit transformations for values bounded in the range [*a, b*]. If transformations are utilized, the perturbation step takes place in the transformed domain [49]. First, the weighted covariance matrix of the **transformed** particles is calculated. Then, a Gaussian step is applied to the transformed particle, and the proposed particle is transformed back to the true domain for the calculation of its likelihood. This transformed random walk breaks the symmetry of the proposal distribution, and the MH ratio must be updated to account for this. This correction comes in the form of the ratio of the transformation Jacobian determinants. For a transformation *ξ*(*θ*_*i*_) *→* Θ_*i*_, let *J*_*ij*_(*θ*_*i*_) = *∂ξ*_*i*_*/∂*(*θ*_*i*_)_*j*_ be the Jacobian of the transformation. Note, for the log and logit transformations, *J* is diagonal. The updated acceptance probability is then

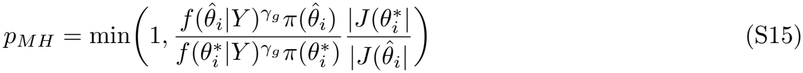

All methods implemented in the <monospace>SMC_Sampler</monospace> class were validated using toy models with conjugate priors or models where the posterior could be directly numerically integrated.

#### Algorithm 1

Sequential Monte Carlo Sampling

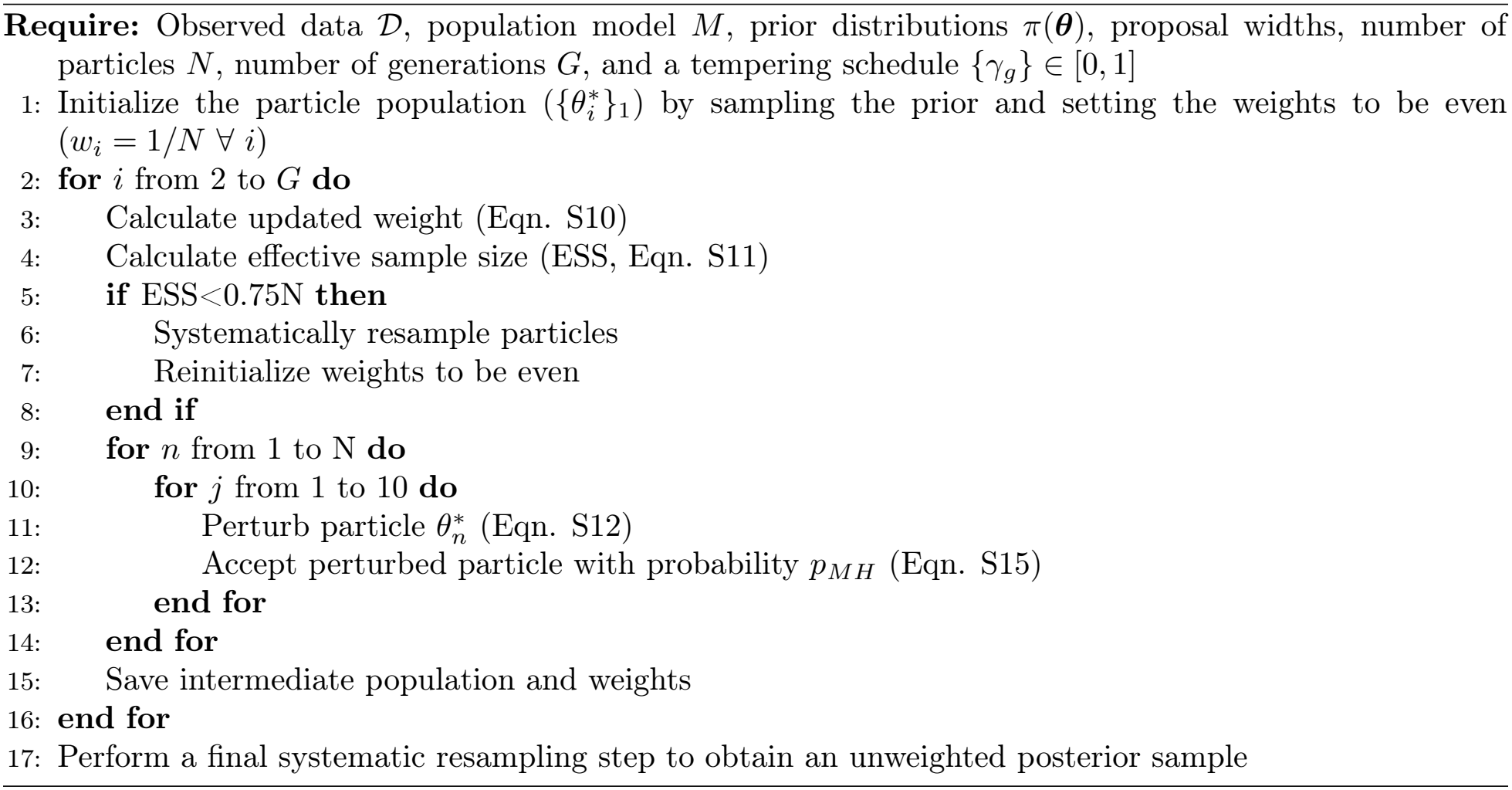

### Detailed Methods for Bayesian Calibration of Hydrogel Population Model Parameters

In this paper, we outline a hierarchical generative statistical framework to capture the natural variability of biologically-derived hydrogels as they cross-link. The deterministic cross-linking relation (Eqn. 4) describes the time evolution of each specimen, while the specimen-level parameters (*y*_0,*i*_, Δ*y*_*i*_, and *τ*_*i*_) are treated as random variables drawn from population distributions. The parameters of this population model were then calibrated in a Bayesian framework using the Sequential Monte Carlo (and Synthetic Likelihood) methods outlined above and the developed MATLAB library. Below, we summarize the specific implementations, including any data processing steps, prior selection, and algorithm settings for each of the three datasets.

#### Shared Implementation Settings

The following settings and priors were used for all three datasets. First, 5,000 particles were used to estimate the posterior over 10 generations, with the values of *γ*_*g*_ ranging linearly from 0 to 1. Log transformations were used for both standard deviation parameters and time constant distribution parameters, and logit transformations were used for the correlation coefficient (bounded [−1,1]) and all effect parameters (bounded [−0.9999, 5]). The prior for the correlation coefficient was set as a Uniform (*U*) distribution over the full range of possible values ([*−*1, 1]), thus being fully uninformative. For all effect parameters, the prior was set as a truncated Normal distribution (*N*_*T r*_), with a location value *µ* = 0 (corresponding to a mode of 0, no change), a scale factor of 1, and a truncation range of [*−*0.9999, 5]. This prior put the most probable values around the null effect and contained 95% of possible effect sizes between approximate −1 and 2. The lower limit of −0.9999 enforced that the temperature would not invert the direction of the sample dynamics (e.g., changing a stiffening to a softening), but was not set to exactly −1, as this was found to cause numerical issues. The upper limit of 5 was set for compatibility with a logit transform which needs an upper limit. Finally, the priors for the time constant parameters (*µ*_*τ*_ and *σ*_*τ*_) were constant across the different data types, as the dynamics of these processes are driven by similar molecular mechanisms over a similar time course. For both values, we used Gamma (Γ) distributions to enforce strict positivity. For the mean time constant, the Gamma distribution (Γ(3, 40)) had a mean of 120 minutes and a standard deviation of about 70 minutes. Prior experiments in the lab have shown that these materials cross-link within ~ 24 hours to the point of forming functional actuators and end effectors. We thus bound the time constant terms to be on the order of a few hours. For the standard deviation of the time constants, the Gamma (Γ(2, 30)) had a mean of 60 minutes and a standard deviation of about 42 minutes. This again reflects previous experience in the lab where the cross-linking time would not vary by more than a couple of hours.

**Table S5:**
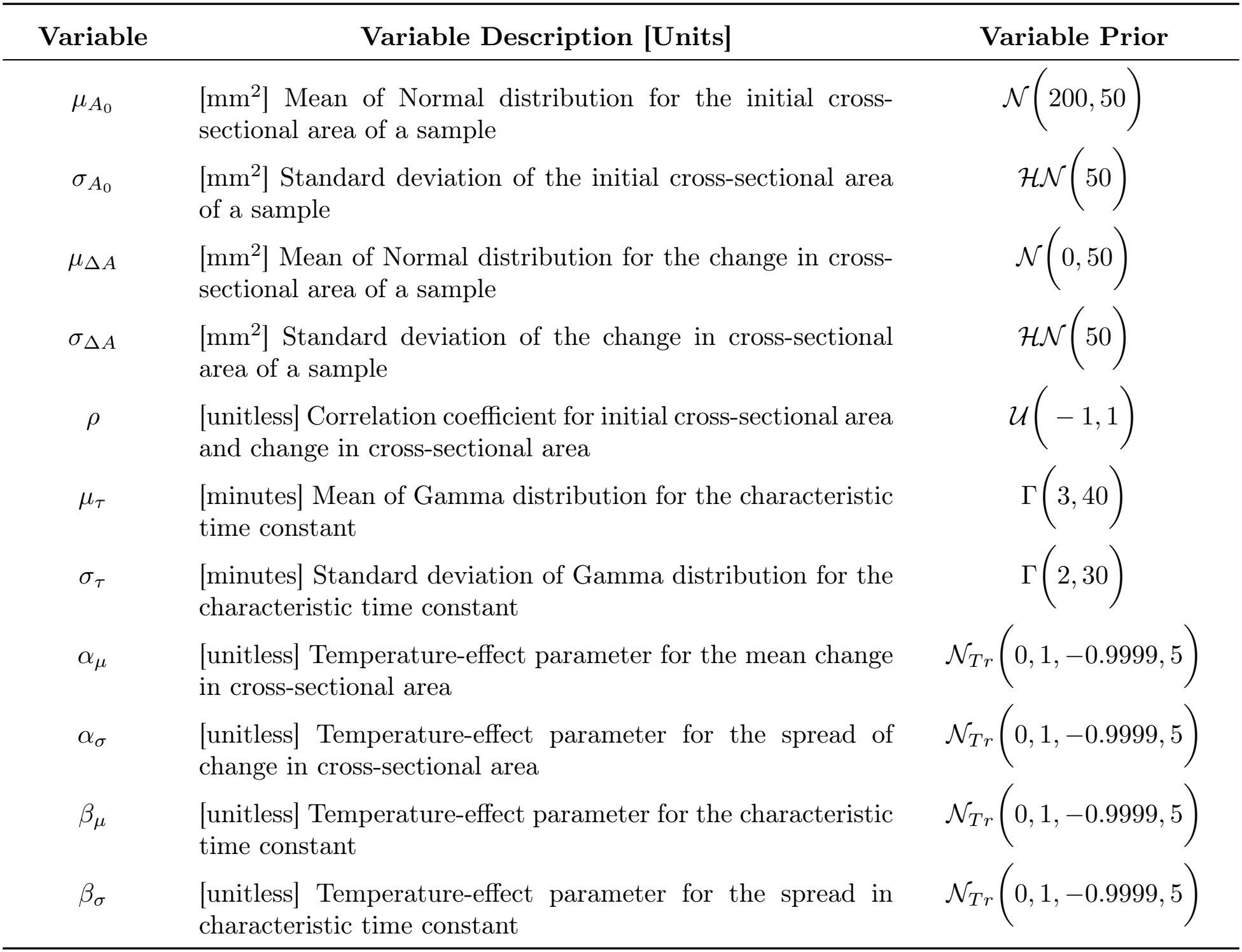
Variables and priors for SMC fitting of the CSA population model.

| **Variable** | **Variable Description [Units]** | **Variable Prior** |
|---|---|---|
| μ<sub>A<sub>0</sub></sub> | [mm<sup>2</sup>] Mean of Normal distribution for the initial cross-sectional area of a sample | 𝒩(200, 50) |
| σ<sub>A<sub>0</sub></sub> | [mm<sup>2</sup>] Standard deviation of the initial cross-sectional area of a sample | ℋ𝒩(50) |
| μ<sub>ΔA</sub> | [mm<sup>2</sup>] Mean of Normal distribution for the change in cross-sectional area of a sample | 𝒩(0, 50) |
| σ<sub>ΔA</sub> | [mm<sup>2</sup>] Standard deviation of the change in cross-sectional area of a sample | ℋ𝒩(50) |
| ρ | [unitless] Correlation coefficient for initial cross-sectional area and change in cross-sectional area | 𝒰( − 1, 1) |
| μ<sub>τ</sub> | [minutes] Mean of Gamma distribution for the characteristic time constant | Γ(3, 40) |
| σ<sub>τ</sub> | [minutes] Standard deviation of Gamma distribution for the characteristic time constant | Γ(2, 30) |
| α<sub>μ</sub> | [unitless] Temperature-effect parameter for the mean change in cross-sectional area | 𝒩<sub>Tr</sub>(0, 1, −0.9999, 5) |
| α<sub>σ</sub> | [unitless] Temperature-effect parameter for the spread of change in cross-sectional area | 𝒩<sub>Tr</sub>(0, 1, −0.9999, 5) |
| β<sub>μ</sub> | [unitless] Temperature-effect parameter for the characteristic time constant | 𝒩<sub>Tr</sub>(0, 1, −0.9999, 5) |
| β<sub>σ</sub> | [unitless] Temperature-effect parameter for the spread in characteristic time constant | 𝒩<sub>Tr</sub>(0, 1, −0.9999, 5) |

#### Algorithm Implementation for Cross-Sectional Area

The population parameters describing the cross-sectional area (CSA) dynamics were fit using the Sequential Monte Carlo workflow implemented in <monospace>AreaDynamicsPopulationModel_SMC.m.</monospace> For this analysis, fitted CSA trajectories were first summarized into sample-level parameters and then used as observations for population inference. To do so, the individual trajectories were fit to Eqn. 4 to obtain sample-level values of *A*_0,*i*_, Δ*A*_*i*_, and *τ*_*i*_. These values were then used to directly calculate the likelihood function throughout the SMC algorithm. The weighted covariance scaling factor *η* was set to 2.38 (corresponding to the theoretical optimal value for random-walk Metropolis-Hastings [48]). Table S5 summarizes the CSA-specific population parameters, their descriptions, and the priors used in posterior estimation. The prior for the mean starting CSA (*µ*_*A*0_) was set as a Normal (*N*) distribution centered on the CSA specific in the CAD (200 mm^2^) with a standard deviation of 50. The mean change in CSA (*µ*_Δ*A*_ was also set as a Normal distribution with standard deviation of 50, but this time centered on 0, reflecting the possibility of either swelling or contracting during cross-linking. Finally, the prior for both standard deviations terms was set as a Half Normal (*ℋN*) distribution with a scale factor of 50, reflecting a broad uncertainty in this variability, but requiring the value to be positive. Figure S4 shows the joint distributions and the convergence of model parameters after 10 generations of SMC.

**Figure S4:**
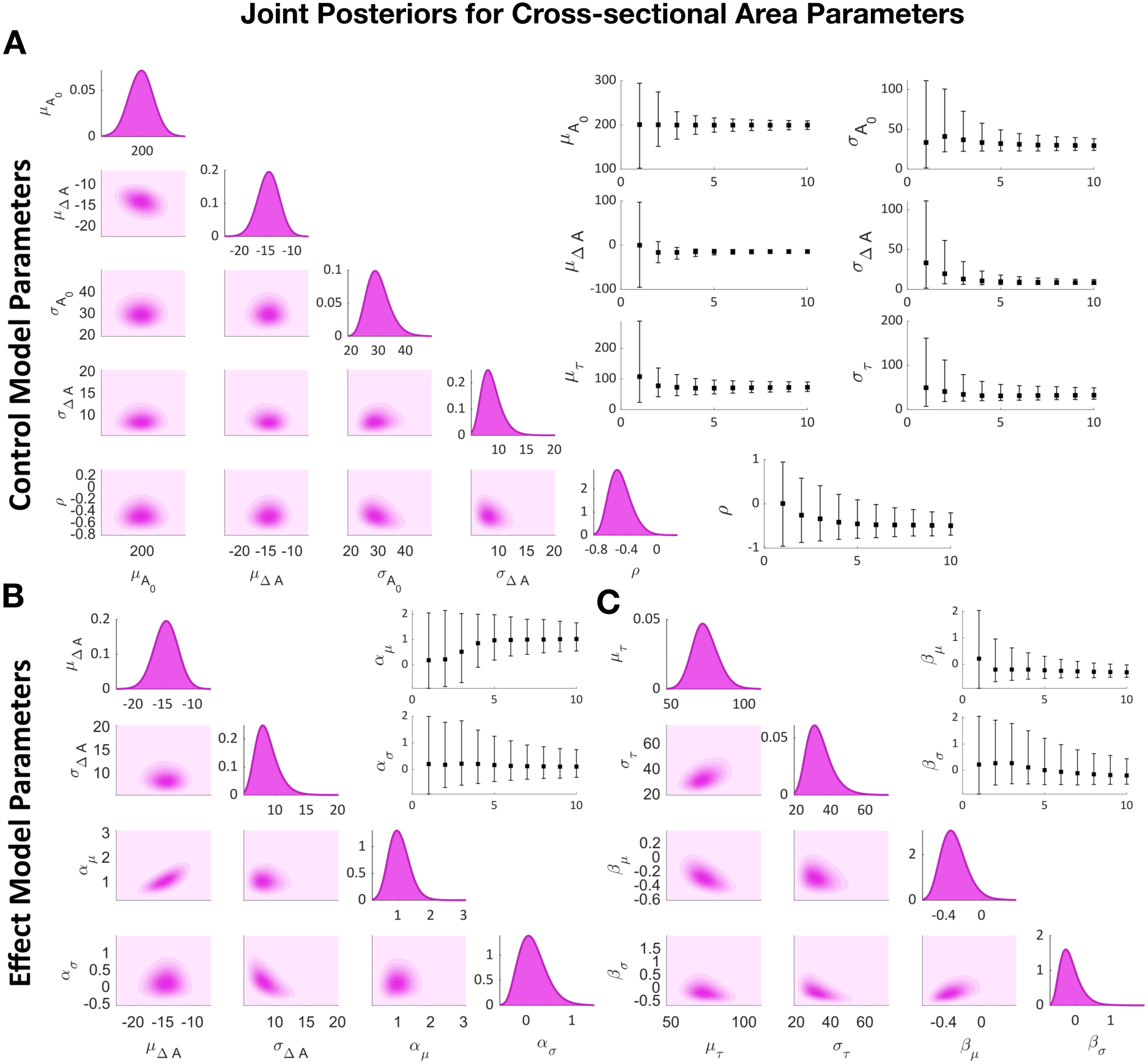
Marginal posterior distributions (diagonals) and joint probability density contours (lower off-diagonals) derived from Sequential Monte Carlo (SMC) sampling, partitioned into three parameter-linked sub-panels for readability. (A) Cross-sectional Area Trajectory Profile (5×5 grid): Joint relationships governing the baseline area 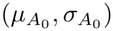, its internal correlation (*ρ*), and the magnitude of area change (*µ*_Δ*A*_, *σ*_Δ*A*_). (B) Δ*A* Change & Temperature Effects (4×4 grid): Direct evaluation of baseline trajectory shifts alongside their corresponding temperature conditional effect terms (*α*_*µ*_, *α*_*σ*_). (C) *τ* Rate & Temperature Effects (4×4 grid): cross-linking rate dynamics and population heterogeneity (*µ*_*τ*_, *σ*_*τ*_) mapped against temperature conditional effect terms (*β*_*µ*_, *β*_*σ*_). For all panels, diagonal curves represent smooth, posterior probability density, while off-diagonal plots display joint probability density contours mapping the highest density regions of the parameter space, with concentric isolines indicating steps of decreasing posterior probability away from the mode. Upper triangles show the median and 95% Credible Interval of parameters over 10 generations of SMC convergence.

#### Algorithm Implementation for Effective Elastic Modulus

Effective elastic modulus was fit using the population-model workflow implemented in <monospace>StiffnessDynamicsPo</monospace> <monospace>pulationModel_ABC_SMC.m</monospace>. No pre-processing was needed for the stiffness values (after calculating from the strain-strain curves), as we cannot fit individual dynamics curves to these data. Instead, the stiffness value at a given time and temperature were directly use in a synthetic likelihood to perform the SMC calibration of the posterior. For these data, the synthetic likelihood was calculated using a univariate Gaussian kernel density estimate over simulated data for a given time and temperature to match the experimental data. The weighted covariance scaling factor was set to 2, as slightly narrower proposals were needed to maintain a good MCMC acceptance rate. Table S6 summarizes the effective elastic modulus population parameters, their descriptions, and the priors used. As existing data for these materials is not available, we utilized fairly uninformative priors, as to not bias the fitting towards a particular region of the parameter space. However, some general bounds could be established. For the stiffness terms, from prior experience with these materials, we knew them to be much softer than typical elastomers (*E ~ O*(10^3^)kPa) and to be quite variable. We thus chose a Normal prior centered the stiffness parameters around a stiffness of ~ 100 kPa, and kept the standard deviations broad. As with the CSA, we centered the mean change on 0 to allow for positive or negative changes. The standard deviation priors were again set to Half-Normal distributions with a broad scale factor. Figure S5 shows the joint distributions and convergence of model parameters after 10 generations of SMC.

**Table S6:**
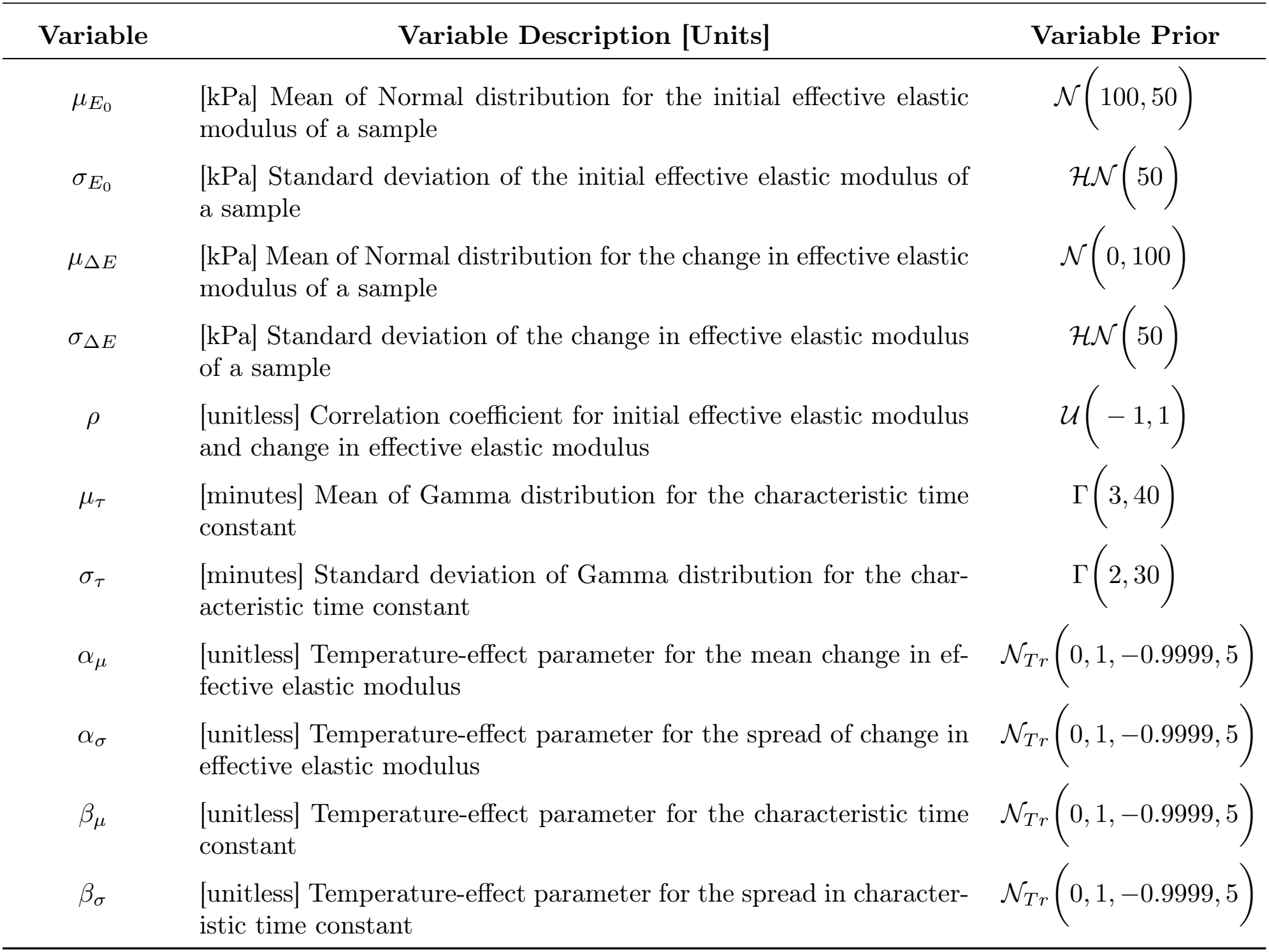
Variables and priors for population-model fitting of the effective elastic modulus data.

| **Variable** | **Variable Description [Units]** | **Variable Prior** |
|---|---|---|
| *μ*<sub>*E*<sub>0</sub></sub> | [kPa] Mean of Normal distribution for the initial effective elastic modulus of a sample | $\mathcal{N}\left(100, 50\right)$ |
| *σ*<sub>*E*<sub>0</sub></sub> | [kPa] Standard deviation of the initial effective elastic modulus of a sample | $\mathcal{HN}\left(50\right)$ |
| *μ*<sub>Δ*E*</sub> | [kPa] Mean of Normal distribution for the change in effective elastic modulus of a sample | $\mathcal{N}\left(0, 100\right)$ |
| *σ*<sub>Δ*E*</sub> | [kPa] Standard deviation of the change in effective elastic modulus of a sample | $\mathcal{HN}\left(50\right)$ |
| *ρ* | [unitless] Correlation coefficient for initial effective elastic modulus and change in effective elastic modulus | $\mathcal{U}\left(-1, 1\right)$ |
| *μ*<sub>*τ*</sub> | [minutes] Mean of Gamma distribution for the characteristic time constant | Γ(3, 40) |
| *σ*<sub>*τ*</sub> | [minutes] Standard deviation of Gamma distribution for the char-acteristic time constant | Γ(2, 30) |
| *α*<sub>*μ*</sub> | [unitless] Temperature-effect parameter for the mean change in ef-fective elastic modulus | $\mathcal{N}_{Tr}\left(0, 1, -0.9999, 5\right)$ |
| *α*<sub>*σ*</sub> | [unitless] Temperature-effect parameter for the spread of change in effective elastic modulus | $\mathcal{N}_{Tr}\left(0, 1, -0.9999, 5\right)$ |
| *β*<sub>*μ*</sub> | [unitless] Temperature-effect parameter for the characteristic time constant | $\mathcal{N}_{Tr}\left(0, 1, -0.9999, 5\right)$ |
| *β*<sub>*σ*</sub> | [unitless] Temperature-effect parameter for the spread in character-istic time constant | $\mathcal{N}_{Tr}\left(0, 1, -0.9999, 5\right)$ |

#### Algorithm Implementation for Mass Reduction

Mass reduction was fit using the population-model workflow implemented in <monospace>MassDynamicsPopulationMod el_ABC_SMC.m</monospace>. As with the stiffness data, no pre-processing was possible. However, as we had both starting and final mass values, the synthetic likelihood was calculated using a bivariate Gaussian kernel density estimate over jointly sampled starting and final masses for a given cross-linking time and temperature. The weighted covariance scaling factor was again set to 2 to slightly narrow the proposal distributions. Table S7 summarizes the mass-reduction population parameters, their descriptions, and the priors used. As with the CSA data, the mean starting mass (Normal prior) was centered on the mass prescribed in the CAD model of the samples, and the change (Normal prior) was centered on 0 (no mass change). The standard deviation priors were set to be broad and strictly positive. Figure S6 shows the joint distributions and convergence of model parameters after 10 generations of SMC.

**Figure S5:**
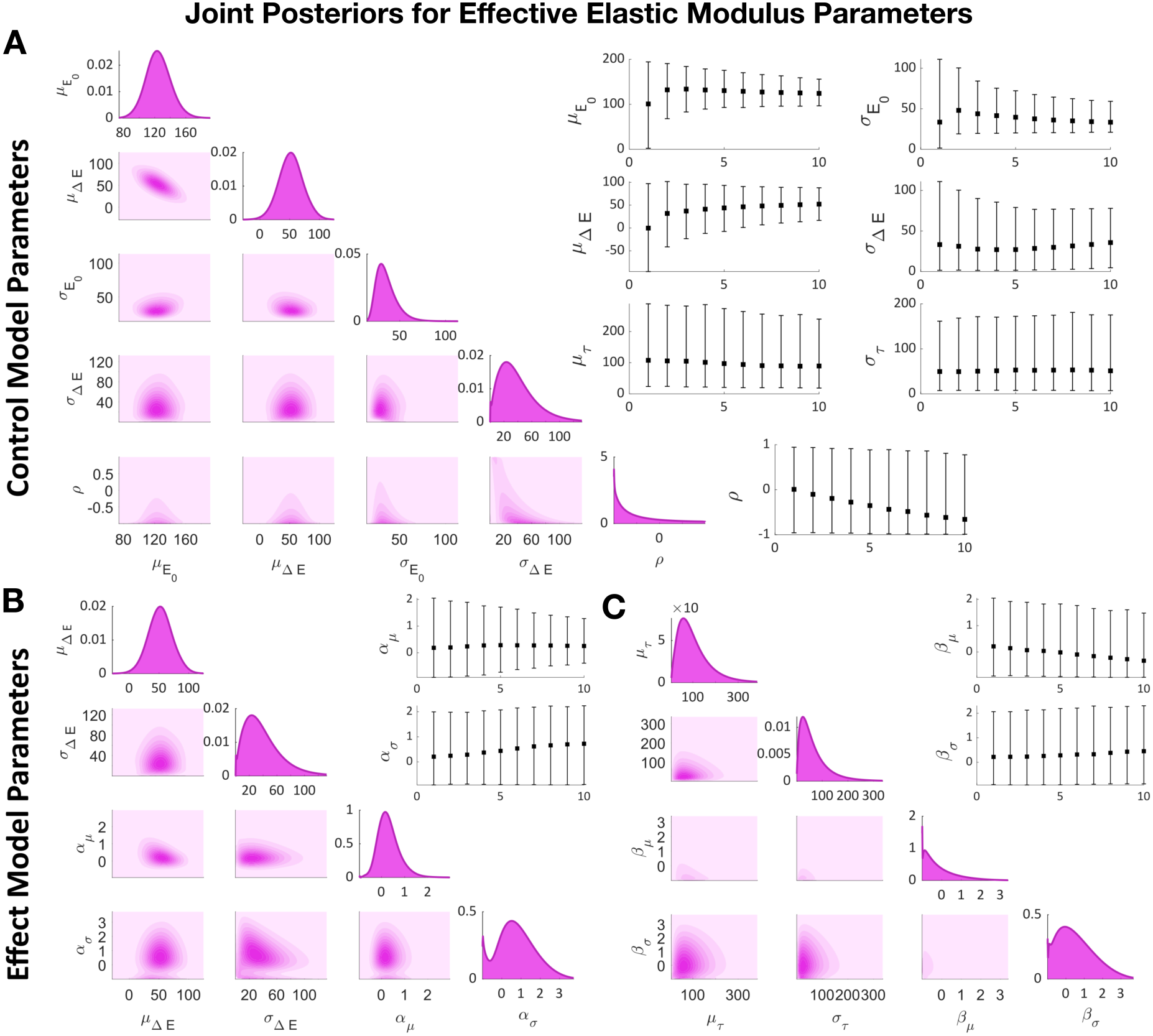
Marginal posterior distributions (diagonals) and joint probability density contours (lower off-diagonals) derived from Sequential Monte Carlo (SMC) sampling, partitioned into three parameter-linked sub-panels for readability. (A) Effective Elastic Modulus Trajectory Profile (5×5 grid): Joint relationships governing the baseline stiffness 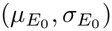, its internal correlation (*ρ*), and the magnitude of stiffness change (*µ*_Δ*E*_, *σ*_Δ*E*_). (B) Δ*E* Change & Temperature Effects (4×4 grid): Direct evaluation of baseline trajectory shifts alongside their corresponding temperature conditional effect terms (*α*_*µ*_, *α*_*σ*_). (C) *τ* Rate & Temperature Effects (4×4 grid): cross-linking rate dynamics and population heterogeneity (*µ*_*τ*_, *σ*_*τ*_) mapped against temperature conditional effect terms (*β*_*µ*_, *β*_*σ*_). For all panels, diagonal curves represent smooth, posterior probability density, while off-diagonal plots display joint probability density contours mapping the highest density regions of the parameter space, with concentric isolines indicating steps of decreasing posterior probability away from the mode. Upper triangles show the median and 95% Credible Interval of parameters over 10 generations of SMC convergence..

**Table S7:**
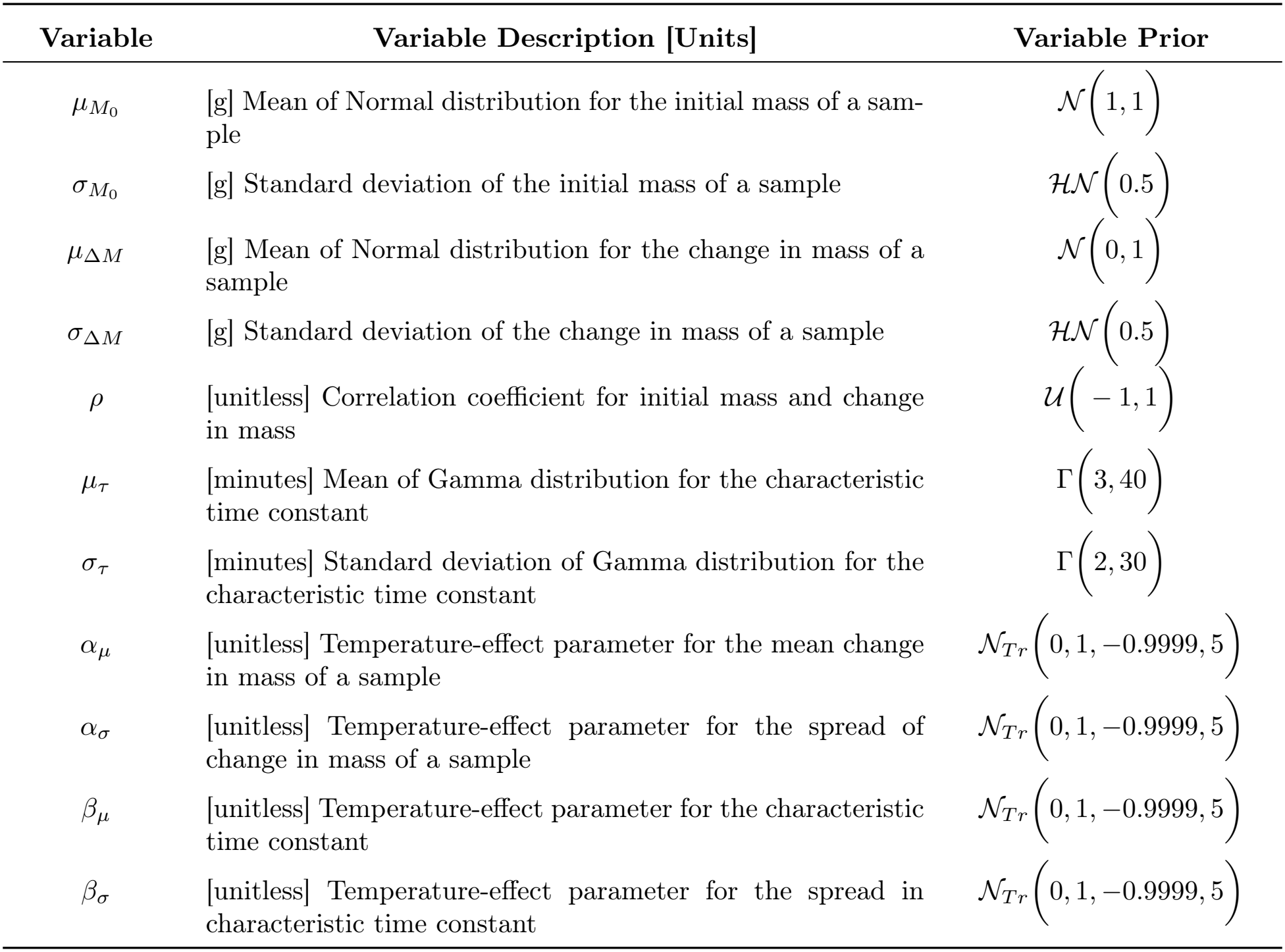
Variables and priors for SMC fitting of the mass reduction population model.

| Variable | Variable Description [Units] | Variable Prior |
|---|---|---|
| μ<sub>M<sub>0</sub></sub> | [g] Mean of Normal distribution for the initial mass of a sam-ple | 𝒩(1, 1) |
| σ<sub>M<sub>0</sub></sub> | [g] Standard deviation of the initial mass of a sample | ℋ𝒩(0.5) |
| μ<sub>ΔM</sub> | [g] Mean of Normal distribution for the change in mass of a sample | 𝒩(0, 1) |
| σ<sub>ΔM</sub> | [g] Standard deviation of the change in mass of a sample | ℋ𝒩(0.5) |
| ρ | [unitless] Correlation coefficient for initial mass and change in mass | 𝒰( − 1, 1) |
| μ<sub>τ</sub> | [minutes] Mean of Gamma distribution for the characteristic time constant | Γ(3, 40) |
| σ<sub>τ</sub> | [minutes] Standard deviation of Gamma distribution for the characteristic time constant | Γ(2, 30) |
| α<sub>μ</sub> | [unitless] Temperature-effect parameter for the mean change in mass of a sample | 𝒩<sub>Tr</sub>(0, 1, −0.9999, 5) |
| α<sub>σ</sub> | [unitless] Temperature-effect parameter for the spread of change in mass of a sample | 𝒩<sub>Tr</sub>(0, 1, −0.9999, 5) |
| β<sub>μ</sub> | [unitless] Temperature-effect parameter for the characteristic time constant | 𝒩<sub>Tr</sub>(0, 1, −0.9999, 5) |
| β<sub>σ</sub> | [unitless] Temperature-effect parameter for the spread in characteristic time constant | 𝒩<sub>Tr</sub>(0, 1, −0.9999, 5) |

**Figure S6:**
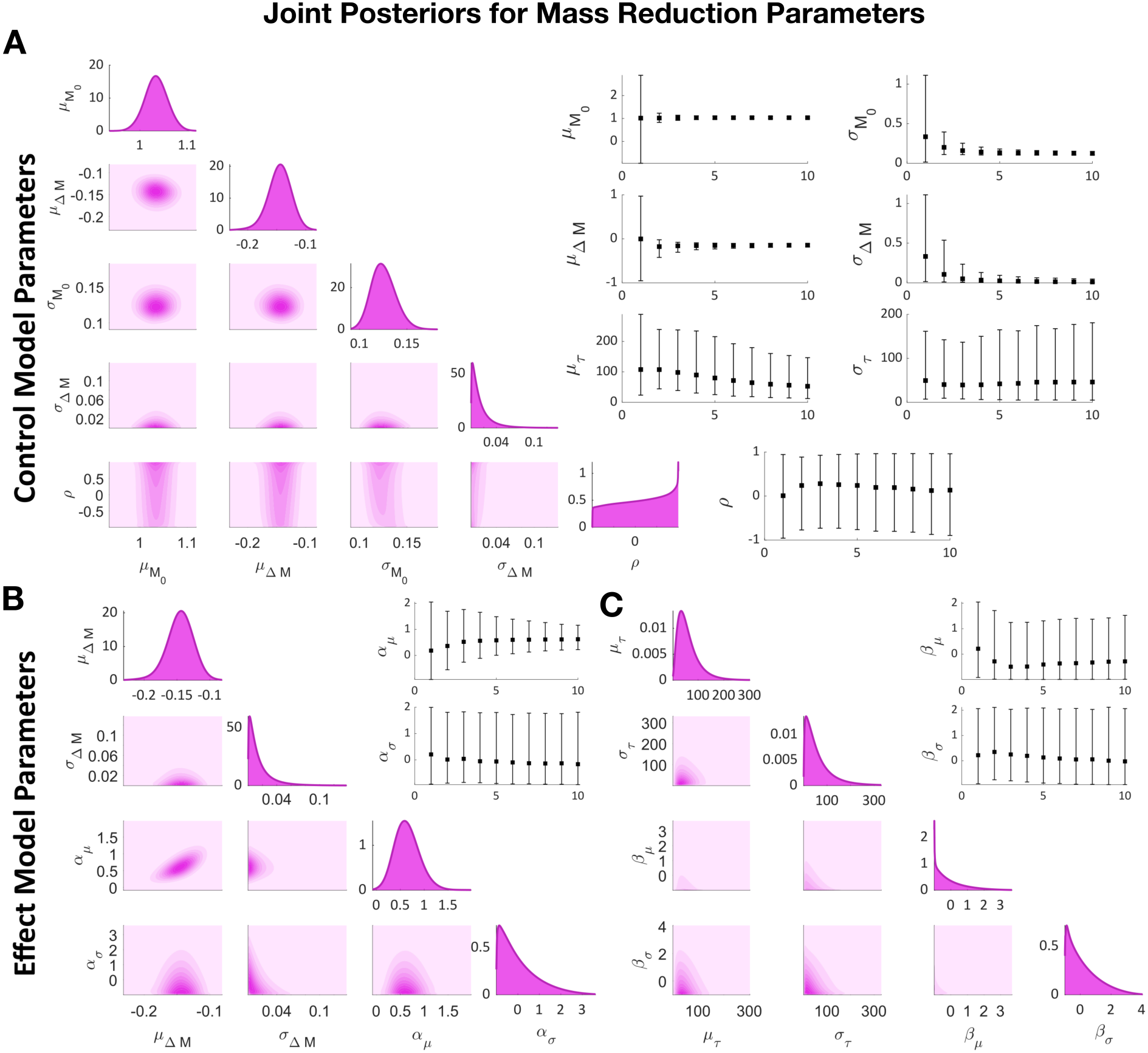
Marginal posterior distributions (diagonals) and joint probability density contours (lower off-diagonals) derived from Sequential Monte Carlo (SMC) sampling, partitioned into three parameter-linked sub-panels for readability. (A) Mass Reduction Trajectory Profile (5×5 grid): Joint relationships governing the baseline mass 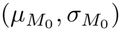, its internal correlation (*ρ*), and the magnitude of mass change (*µ*_Δ*M*_, *σ*_Δ*M*_). (B) Δ*M* Change & Temperature Effects (4×4 grid): Direct evaluation of baseline trajectory shifts alongside their corresponding temperature conditional effect terms (*α*_*µ*_, *α*_*σ*_). (C) *τ* Rate & Temperature Effects (4×4 grid): cross-linking rate dynamics and population heterogeneity (*µ*_*τ*_, *σ*_*τ*_) mapped against temperature conditional effect terms (*β*_*µ*_, *β*_*σ*_). For all panels, diagonal curves represent smooth, posterior probability density, while off-diagonal plots display joint probability density contours mapping the highest density regions of the parameter space, with concentric isolines indicating steps of decreasing posterior probability away from the mode. Upper triangles show the median and 95% Credible Interval of parameters over 10 generations of SMC convergence.

